# Engineering Stable Hydrogels with Polydisperse Yeast Exopolysaccharides for Embedding Cancer Spheroids

**DOI:** 10.64898/2026.02.04.703759

**Authors:** Henrique Sepúlveda Del Rio Hamacek, Tobias Butelmann, Katharina Ostertag, Kerit-Lii Joasoon, Oksana Tingajeva, Piia Jõul, Petri-Jaan Lahtvee, V. Prasad Shastri, Rahul Kumar

## Abstract

Polysaccharides are often used to mimic physiological environments such as for cancer research models. However, established polysaccharides can display limited long-term stability and high batch-to-batch variability. To overcome this, biomanufactured polysaccharides are increasingly utilized in biomaterials. Here, we produced and characterized *Rhodotorula toruloides* yeast exopolysaccharides (EPS) and used it to engineer hydrogel for culturing cancer cells. Yeast fermentation of glucose, mannose, and xylose yielded varying EPS amounts (1.68, 1.44, and 0.48 g/L, respectively) with similar compositions, suggesting a common biosynthetic pathway. The glucose-derived EPS characterization identified multiple linkage types and three molecular weight fractions (1.75, 30.0, and 1000 kDa), and its solutions exhibited Newtonian behavior, indicating minimal chain-chain interactions. Solubilizing this polydisperse EPS with polyethylene glycol diacrylate and UV-crosslinking it enabled the engineering of semi-interpenetrating polymer network hydrogel that efficiently embedded cancer spheroids. Our study introduces an integrated biomanufacturing strategy to generate stable and consistent biomaterials, applicable for tissue engineering.

**Graphical abstract:** 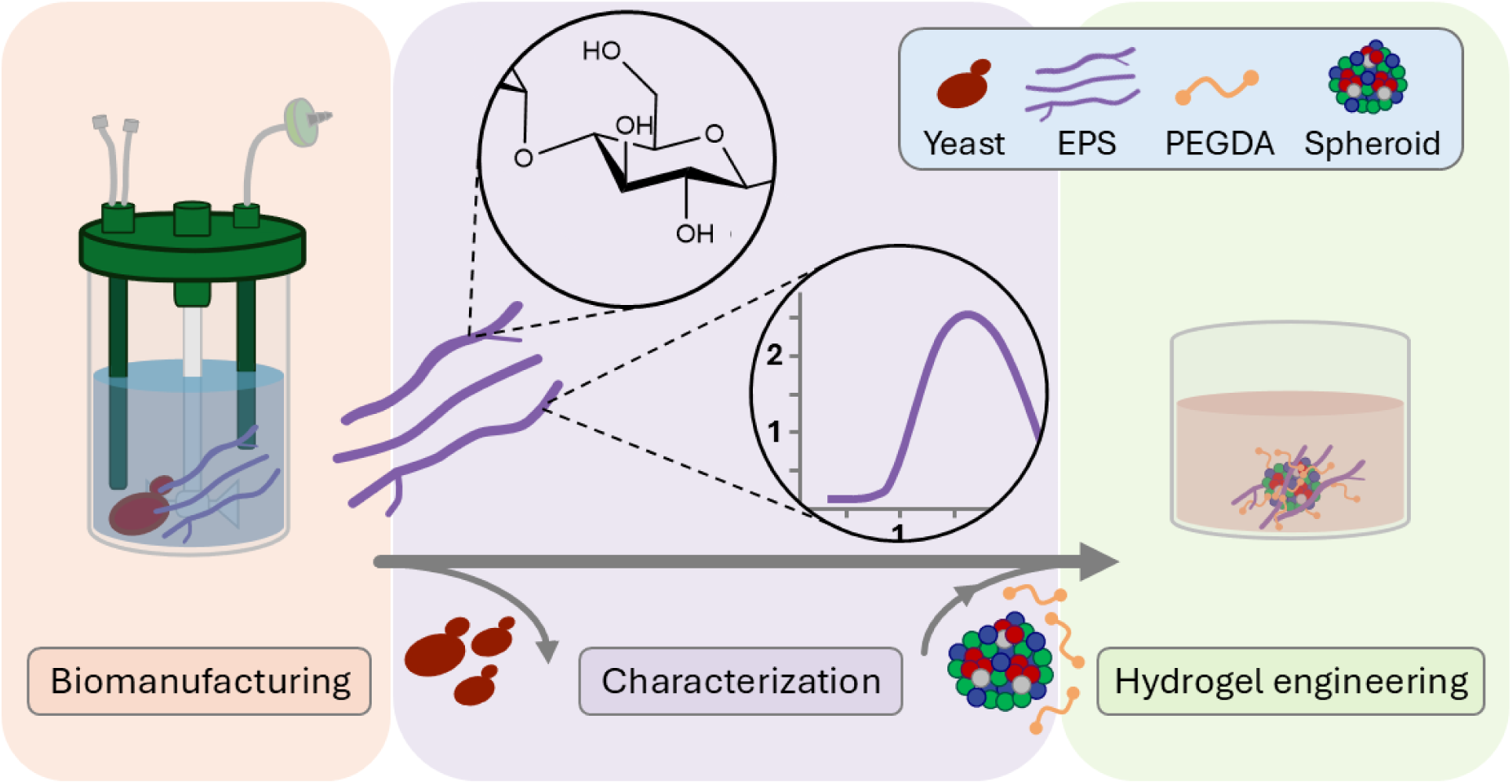

## 1. Introduction

Most industrially relevant polysaccharides are currently extracted from natural resources, such as alginate from seaweeds, chitin from crustaceans, and cellulose from plants, whereas biomanufactured polysaccharides produced by living cells^1^, such as exopolysaccharides (EPS), remain underrepresented^2^. However, when compared to nature-sourced polysaccharides, microbial EPS have a great potential in addressing problems surrounding quality, consistency, and diversity in physicochemical characteristics, especially for niche applications, such as cosmeceuticals^3^. For instance, while the quality and properties of marine polysaccharides depend on seasonal variables and are impacted by environmental factors, such as water temperature, salinity, and a more complex extraction process, the biomanufacturing of EPS can be carried out in a controlled bioreactor, and their composition precisely engineered using genetic tools^4–7^.

Prominent examples of microbial EPS produced at large scales for food-related applications are xanthan, gellan, dextran, pullulan, and curdlan^8^. For biomaterial applications, the use of EPS depends on their structural, physicochemical, and biological properties, which are typically influenced by their native functions for the producer microbes^9,10^. Consequently, screening of potential EPS producing microbes, their physiology, and characterization of EPS remain vital for broadening the repertoire of available microbial cell factories employed in biomanufacturing^11,12^.

Specifically, tissue engineering, regenerative medicine, drug delivery, and 3D cancer cell cultures applications widely utilize natural polysaccharides to generate hydrogel-based biomaterials for cells encapsulation^13,14^, often by combining them with synthetic polymers, such as polyethylene glycol diacrylate (PEGDA)^15–17^. This blending creates a highly controllable and physiologically relevant environment for cell cultures by diminishing the limiting attributes of each type of polymer and imparting beneficial characteristics to the engineered hydrogels^18,19^.

Pertinent to the above discussion, in our previous research, we identified the physiological mechanism by which the oleaginous yeast *Rhodotorula toruloides* produces EPS^20^. However, it remains unexplored how different sugars affect EPS production and composition in *R. toruloides*, what the resulting EPS rheological properties are, and how this EPS could be leveraged for biomaterial applications. Hence, this study aimed to address these previously unexamined aspects of the *R. toruloides* biosynthesized EPS and to demonstrate its use by combining it with PEGDA to form a semi-interpenetrating polymer network (semi-IPN). We hypothesized that this EPS-PEGDA semi-IPN could support stable mammalian spheroid cultures under physiological conditions. In conclusion, our study presents an integrated biomanufacturing strategy for the production, characterization, hydrogel engineering of a yeast-derived EPS, and use of it for embedding and culturing cancer spheroids, indicating a path for sustainable materials sourcing and utilization in biotechnology and biomedicine.

## 2. Materials and Methods

### 2.1. Yeast strain and culture conditions

*Rhodotorula toruloides* (genotype - CCT0783, NBRC10076) yeast was used throughout this study^20^. The glycerol stocks were stored at -80 °C and thawed on ice prior to inoculation. Pre-culture cultivations were performed in a centrifuge tube (50 mL) or a shake flask (250 mL), containing 5 mL or 25 mL of working volume of YPD, respectively. The pre-culture was incubated overnight (200 rpm, 30 °C) in a rotary shaker incubator (Excella E25, New Brunswick Scientific, USA) and used as the inoculum source for experiments with chemically defined media. Before inoculating suspension cultures, the optical density at 600 nm (OD_600_) was measured to estimate the inoculum volume needed to achieve an OD_600_ equal to 0.1, unless stated otherwise. Suspension cultivation experiments were performed using pre-sterilized 250-mL shake flasks with 25 mL of chemically defined media at 200 rpm. The temperature was kept constant at 30 °C. The cultivations were carried out for 96 h. Samples for dry cell weight (DCW), EPS, and pH measurements were collected every 24 h. All culture preparations and samplings were performed under sterile conditions.

### 2.2. Culture media composition and preparation

Culture media, YPD or a chemically defined composition, were prepared according to previous protocols^21^. The YPD medium contained 20 g of peptone (bacteriological grade, >95%, Thermo Fisher Scientific, USA), 10 g of yeast extract (Thermo Fisher Scientific, USA), and 22 g of D(+)-glucose monohydrate. For the chemically defined medium, D(+)-glucose monohydrate (>99.5%, Carl Roth GmbH, Germany), D(+)-mannose (≥99%, Thermo Fisher Scientific, USA), D(+)-xylose (>99.5%, Carl Roth GmbH, Germany), or D(+)-galactose (Naxo, Estonia) was used as the carbon source to give a concentration of sugar equal to 20 g L^-1^. 5 g of (NH_4_)_2_SO_4_ (99.8%, Thermo Fisher Scientific, USA), 3 g of KH_2_PO_4_ (100%, Thermo Fisher Scientific, USA), and 0.5 g of MgSO_4_•7H_2_O (≥99%, Thermo Fisher Scientific, USA) were dissolved together in ultrapure water. Thermal sterilization was performed in an autoclave (Systec V-95, Systec, USA) at 121 °C for 15 minutes (2 bar). The carbon source solution was sterilized separately and, after cooling down, was added to prepare 1 L of medium, supplemented with 1 mL each of vitamins and trace elements from stock solutions. The composition used to prepare the stock solutions was previously described^21^. The pH of prepared media was not adjusted and was approximately 4.50 at the start of the cultivation.

### 2.3. Bioreactor cultivations

For a bioreactor cultivation of *R. toruloides*, method and conditions described in the literature were adapted^22^. Briefly, 1 L stirred tank bioreactors MiniBio 1000TM (Applikon Biotechnology, Netherlands) were used in batch mode. The working volume in the bioreactors was 0.9 L, containing chemically defined media with a glucose concentration of 40 g L^-1^. The other components and nutrients of the medium were kept at the same concentrations described in previous sections. Antifoam 204 (Sigma-Aldrich, USA) was added at 0.1 mL per liter of liquid culture to stop foam formation. The stirring speed and airflow in the tanks were set to 750 rpm and 700 mL air/min, respectively. The temperature was kept constant at 30 °C. The pH was monitored throughout cultivation with a Sensor pH Gel 185/8 Minibio 100 (Applikon Biotechnology, Netherlands). The O_2_ and CO_2_ concentrations of the outflow gas were obtained using off-gas sensors BlueInOne (BlueSens, Germany). The data collection was done by Lucullus PIMS Lite v3.7.4 software (Getinge, Sweden). All bioreactor experiments were performed in triplicate.

The pre-inoculum was obtained as previously described. To inoculate the bioreactors, the cells from the pre-inoculum were centrifuged (5 min, 6000 rpm, 20 °C) and resuspended in chemically defined media. Cells were added to the bioreactors to give an initial OD_600_ equal to 0.2. During cultivation, different sample volumes were collected at various time points to measure the OD_600_ (1 mL), DCW (1 mL), and EPS production (5 mL). After 120 h, the cultivation was stopped, and each batch was processed to extract EPS and quantify glycerol by HPLC analysis.

### 2.4. Extraction and quantification of EPS in suspension cultures

Previously established methods for the extraction and quantification of EPS from the suspension cultures were used^20^. Briefly, the culture medium was centrifuged (5000 × *g*, 4 °C, 30 minutes), and the resulting supernatant, which contained the suspended EPS, was carefully separated from the cell pellet. To precipitate the EPS, 96 vol.% ethanol was added to the supernatant at a ratio of two parts in volume of ethanol to one part of supernatant. The mixture was kept at 4 °C for 24 h to promote EPS precipitation. After this period, two distinct layers of EPS formed in the middle/top of the tubes and at the bottom. The top layer was carefully collected with tweezers. For bioreactor experiments, a bottom-layer EPS precipitate was obtained by centrifugation (13000 × *g*, 4 °C, 10 min) and the resulting pellet was collected. Finally, the top and bottom EPS fractions were dried separately in an incubator at 50 °C for 48 to 72 h, until no change in mass was observed, indicating that all the excess liquid evaporated.

### 2.5. Bioprocess parameter calculations

#### 2.5.1. Yields from biomass and substrates

To assess the bioprocess performance, the product yield from biomass (Y_PX_), the biomass yield from substrate (Y_XS_), and the product yield from substrate (Y_PS_) were calculated using Equations 1, 2, and 3, respectively. The product (P) denotes the EPS, biomass (X) is the DCW, and substrate (S) is the amount of substrate consumed. For each reported yield, the units are either grams or mols, the calculation interval is specified, and whether the yield represents an overall (cumulative) or instantaneous value is indicated.

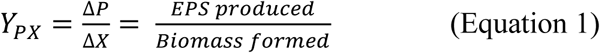

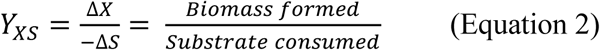

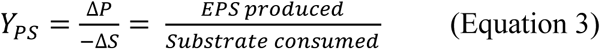

#### 2.5.2. Maximum specific growth rate (μ_max_)

To determine the maximum specific growth rate (μ_max_), which is the specific growth rate (μ) at the exponential growth, it was indirectly calculated by using the CO_2_ production rate (R_CO2_). The identification of the exponential growth period was done manually by looking for linear windows in the natural logarithm of CO_2_ production rate *versus* time plot and taking the region with the highest slope (Equation 4 and 5). For this approximation, the growth rates (μ) and specific CO_2_ production rates (qCO_2_) were assumed to be constant during the exponential phase, and the non-growth associated CO_2_ was negligible.

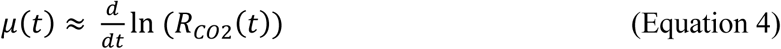

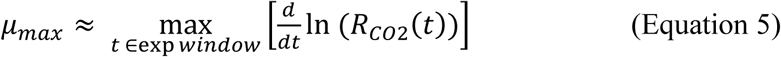

#### 2.5.3. Specific production/consumption rates (r) and respiratory quotient (RQ)

The specific production rate of a product (P) was calculated by dividing the mass or moles per unit time by the mass of the dry cell weight (X, g DCW) obtained in the same time interval (Equation 6). Here, P denotes either EPS or CO_2_. The specific consumption rate of a substrate (S) was calculated analogously (Equation 7) and, in this study, S refers to O_2_. Finally, the respiratory quotient (RQ), which represents the relationship between the produced CO_2_ and the consumed O_2_, was calculated using Equation 8, where n_CO2_ and n_O2_ denote the cumulative amount (mol) of CO_2_ produced and O_2_ consumed in a specified time interval, respectively.

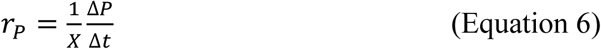

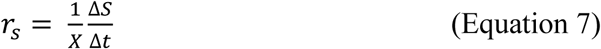

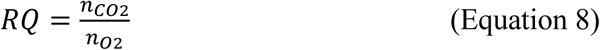

### 2.6. Chemical and Physical Characterizations

#### 2.6.1. High-Performance Liquid Chromatography (HPLC)

For HPLC, samples of 1 mL of medium were collected in 2 mL microcentrifuge tubes, centrifuged (21950 × *g*, 4 °C, 5 min), and the supernatant was subsequently transferred to HPLC vials. The samples were filtered (0.22 μm) and diluted as necessary. The amount of the carbon source (glucose, mannose, galactose, and xylose) and metabolite (glycerol) in different media during cultivation was quantified using HPLC (LC-2030C Plus, Shimadzu, Kyoto, Japan), equipped with a refractive index detector (RID-20A, Shimadzu, Kyoto, Japan). The samples were stored at 4 °C on the sample rack, and 20 μL aliquots were automatically injected onto an Aminex HPX-87H ion exclusion column (Bio-Rad, US). Elution was performed in isocratic mode with a H_2_SO_4_ solution (5 mM) as the mobile phase at a flow rate of 0.6 mL min^-1^ and with column and detector temperatures at 45 °C.

For samples containing EPS, further purification was needed to protect the column integrity. Therefore, to completely remove all macromolecules from the culture medium, 0.2 mL of ethanol (96 vol.%) was added to a 0.1 mL of media sample in a microcentrifuge tube. The tube was vortexed to allow proper mixing, centrifuged (21950 × *g*, 4 °C, 10 min), and 0.1 mL of the supernatant was collected into a new tube. The ethanol mixing and centrifugation procedures were repeated once under the same conditions. The clarified supernatant was carefully transferred with a pipette to an HPLC vial and injected into the machine as described above.

#### 2.6.2. pH measurements

The pH was measured using a HACH HQ30d pH meter (METTLER TOLEDO, USA) from 1 mL samples collected at specific points of cultivation. Samples were prepared by centrifugation (21950 × *g*, 20 °C, 5 min) to remove the cells, and then, the supernatant was collected for the pH measurement.

#### 2.6.3. DCW Determination

The DCW was obtained by collecting 1 mL of cultivation medium and then centrifuging it for 5 minutes at 15200 rpm (21950 × *g*). The supernatant was removed, and 1 mL of saline solution (0.9 wt./vol.% NaCl) was added to resuspend the cells. The samples were vortexed, then pipetted onto pre-weighted 0.45 μm filter papers, and filtered using a vacuum filtration system. While still connected to the vacuum system, thorough washing was done with distilled water to remove residual salts. The filter papers containing biomass were dried in a microwave oven at 900 W for 2-3 min in 30-s intervals. The filter papers were stored in a desiccator until weighed on the following day.

#### 2.6.4. Fourier Transform Infrared (FTIR) Spectroscopy analysis

Dry EPS samples were placed on the attenuated total reflection (ATR) crystal in an FTIR spectrometer, IRTracer-100 (Shimadzu, Japan), and analyzed in ATR mode. The spectra were recorded in the range of 4000 to 400 cm⁻¹, with 6 cm⁻¹ spectral resolution, and with a 5 mm aperture. The peaks identified by the software LabSolutions (Shimadzu, Japan) were compared to literature references to determine the functional groups present in the samples.

#### 2.6.5. EPS compositional analysis

##### 2.6.5.1. Monosaccharide quantification

The monosaccharide composition of the EPS obtained from yeast grown in different carbon sources was determined using previous methods^20^. Briefly, EPS samples (10 mg) were dissolved and hydrolyzed in 1 mL of trifluoroacetic acid (TFA, 2 mol L^-1^) at 120 °C for 90 min in a dry heat block (Digital Dry Bath JXDC-10, Tuohe Electromechanical Technology Co., China). The liquid was evaporated under a nitrogen (N_2_) stream, and 1 mL of sodium borohydride solution (NaBH_4_, 20 g L^-1^) was added. The mixture was allowed to react for 3 h (room temperature) and dried again with N_2_ gas. For derivatization, pyridine (300 μL) and acetic anhydride (400 μL) were added to the vial and heated under mild agitation (90 °C, 30 min) in a dry heat block. Finally, each sample was centrifuged, and the supernatant was transferred to a new glass vial to separate the solid residue.

The compositional analysis was carried out using a 7890A gas chromatograph coupled to a 5975C mass spectrometer (GC-MS, Agilent Technologies, USA) with an electron ionization source and a quadrupole mass analyzer. The sample (1 μL) was injected into the GC-MS using split mode (1:10) at 275 °C. The flow rate of the carrier gas (helium) was 1.3 mL min^-1^. The initial oven temperature was held at 50 °C for 1 min, ramped at 10 °C min^-1^ until it reached 220 °C and held for 5 min, then ramped at 10 °C min^-1^ to 250 °C and held for 2 min. The total running time was 28 min. The analyte ionization was performed in electron ionization mode using an electron energy of 70 eV. The interface, ion source, and mass analyzer temperatures were set to 280, 230, and 150 °C, respectively. Scan mode in the target ion range, from 50 to 800 m/z, was used for monitoring compounds of interest^9,11^. Compounds were separated by a ZB-5plus capillary column (30 m x 0.25 mm x 0.25 μm, Agilent Technologies, USA). Agilent MassHunter Qualitative, Quantitative, and Unknown Analysis software (Version B.07.00, Agilent Technologies, USA) was used for data analysis. The retention times for different sugars were obtained with pure monosaccharide standards that underwent the same derivatization procedure.

##### 2.6.5.2. Glycosidic bond determination

To determine the glycosidic bonds present in the EPS, the methods described by Hamidi et al. and Jiao et al. were slightly adapted^23,24^. In a glass vial, dried EPS (4 mg) was dissolved in DMSO (500 μL, ≥99%). Next, fine NaOH powder (20 mg) was added to each sample, followed by the addition of iodomethane (100 μL, 99.0%, Sigma-Aldrich, USA). The mixture was kept under agitation for 10 min to complete the methylation process. Then, a liquid-liquid extraction was performed with ultrapure water and chloroform (CHCl_3_). CHCl_3_ (1 mL) was added to the reaction mixture and washed with water (1 mL) three times to remove water-soluble compounds. The water layer was removed by pipetting. After the last step, excess anhydrous sodium sulfate (Na_2_SO_4_, >99.0%, Sigma-Aldrich, USA) was added to the CHCl_3_ layer to remove residual water.

Finally, the supernatant of the CHCl_3_ and Na_2_SO_4_ mixture was transferred to a clean glass vial to separate it from the hydrated Na_2_SO_4_. The liquid was dried under a continuous N_2_ stream, resulting in dried methylated EPS. The hydrolysis, derivatization, and GC-MS analysis were performed as described in the previous section by adjusting the reagent quantities to match the methylated polysaccharide weight. To identify glycosidic linkages, Agilent MassHunter Qualitative, Quantitative, and Unknown Analysis software (Version B.07.00) was used to search for partially O-methylated alditol acetates (PMAAs), and available literature was used to match the PMAAs to their respective glycosidic bonds.

##### 2.6.5.3. Protein quantification

Protein quantification was performed with Pierce^TM^ Bradford Protein Assay Reagent following the supplier’s protocol for standard test tube procedures. Briefly, 30 μL of the samples or the standards were added to a cuvette containing 1.5 mL of the Bradford reagent. The cuvettes were incubated in the dark (10 min, room temperature), after which the absorbance was measured at 595 nm. Bovine serum albumin (BSA) stock solution (2 mg mL^-1^) was used as the standard to prepare the calibration curve according to the supplier’s instructions.

#### 2.6.6. Gel permeation chromatography

To determine the molecular weight of the EPS through gel permeation chromatography (GPC), the sample was prepared by dissolving 1.0 mg of dried crude EPS in 1.0 mL of PBS buffer (0.01 M, pH 7.4). Before injection, the solution was filtered through a 0.45 µm poly(ether-sulfone) filter.

The samples were eluted using PBS (0.01 M, pH 7.4) with a flow rate of 1.0 mL min^−1^. The columns used included a precolumn (PSS Suprema Guard, particle size 10 µm, inner diameter: 0.8 cm, length: 5 cm) followed by a series of PSS Suprema columns with a pore size of 100 Å, and then two times 3000 Å (particle size 10 µm, inner diameter: 0.8 cm, length: 30 cm). 50 µL of sample was injected into the system with the autosampler PSS SECurity 1200 (Agilent Technologies, USA) and measured with a refractive index detector 1260 (Agilent Technologies, USA). The system was run at 50 °C and calibrated using pullulan standards from 180 Da to 1.5 MDa. Integration boundaries were 19.5 to 26.5 min, 26.5 to 30.1 min, and 31.35 to 33 min for fractions 1, 2, and 3, respectively. The chromatogram was subsequently analyzed using PSS-WinGPC UniChrom Version 8.33.

#### 2.6.7. Rheological analysis

The apparent viscosity of the culture supernatants containing EPS and EPS solutions at different concentrations (1, 5, 10, 20, and 30 g L^-1^) was measured by adding 15 mL of the sample to a Rheolab QC rheometer (Anton Paar, Austria) equipped with a concentric cylinder measuring system. The shear rate varied from 1 to 100 s^-1^ to give the curve of apparent viscosity^25^. The temperature during the analysis was maintained at 25 °C.

### 2.7. Mammalian cell cultures

#### 2.7.1. Hydrogel preparation, inversion tests, and embedding spheroids

To prepare an EPS solution, EPS (40 mg) was added to Dulbecco’s PBS (DPBS, Gibco, Germany, 1mL) and dissolved overnight at 4 °C to obtain a 4% solution. Then, 40 mg of PEGDA (M_n_ 700 Da, Sigma Aldrich, Germany) and the EPS solution were mixed to prepare 4% EPS + 4% PEGDA (4% EPS-PEGDA) as hydrogels. A 4% PEGDA solution in DPBS was prepared similarly. Next, 2.5 µL of the photo-initiator (2-hydroxy-2-methylpropiophenone, Sigma Aldrich, Germany) was added to 1 mL of the solution. When needed, a spheroid (see preparation in next sections) was added to 4% EPS-PEGDA or 4% PEGDA, and a 30 µL drop of the different solutions was dispensed on a 2.5 x 2 cm glass slide. To test the gelation of the solutions, the slides were tilted perpendicularly to the ground, and the solutions were allowed to flow down the slide for 30 s before and after crosslinking (inversion test). Curing under ultraviolet light (λ = 365 nm) was carried out for 1 min and at a 5 cm lamp distance, using the Inkredible+ bioprinter blue light LED (Cellink, Sweden). Embedded spheroids were cultured at 37 °C under 5% CO_2_ at ∼90% humidity.

#### 2.7.2. Mammalian cell culture

Cells for spheroid preparation were cultured at 37 °C under 5% CO_2_ at ∼90% humidity in their respective medium. For passaging, cells at 70–80% confluency were washed with DPBS and trypsinized (0.05% trypsin/0.02% EDTA, PAN Biotech, Germany) for 5 min or until most of the cells detached. MDA-MB-231 cells were provided by the BIOSS toolbox (Centre for Biological Signaling Studies, University of Freiburg, Germany) and cultured in Dulbecco’s Modified Eagle’s Medium (DMEM) (Gibco, Thermo Fisher, USA) supplemented with 10% fetal bovine serum (FBS) (Thermo Fisher, USA) and 1% penicillin-streptomycin (PAN Biotech, Germany). Human pulmonary microvascular endothelial cells (HPMEC) were acquired from PromoCell (Heidelberg, Germany) and were propagated in a Growth Medium MV (Ready-to-use) (PromoCell: C-22020). Human marrow-derived mesenchymal stem cells (MSC) were kindly provided by Dr. Andrea Barbero and were obtained from patients with informed consent under the regulations of the institution’s ethics committee (University Hospital Basel; ref. nr. of local ethical committee 78/07). They were sub-cultured in alpha-MEM (Thermo Fisher, USA) containing 10% FBS, 1% penicillin-streptomycin, and 5 ng mL^-1^ fibroblast growth factor-2 (R&D Systems, USA).

#### 2.7.3. Lentiviral transduction

Lentiviral particles containing BFP: pLVX-mTagBFP2-P2A-Puro and E2 Crimson: pLVX-E2-Crimson-P2A-Puro (BIOSS Toolbox, University of Freiburg, Germany) were produced in HEK293 cells by mixing transgene vector and packaging vectors (pCMVdR8.74 (packaging plasmid; Addgene, Plasmid #22036) and pMD2.G (envelope plasmid, Addgene, Plasmid #12259)) using branched polyethyleneimine (bPEI) (M_W_ = 25 kDa, Sigma Aldrich, Germany) as the transfection reagent.

For transfection, 5 μg of DNA at a ratio of 4:3:1 (transgene: packaging: envelope plasmids) was diluted in 250 μL of Opti-MEM (Invitrogen, Thermo Fisher, USA), and 11.25 μL of bPEI (1 mg mL^-1^) was added and incubated for 25 min at room temperature before transferring to HEK293 cells. 16 h after transfection, the medium was exchanged with the medium of the target cells. 24 and 48 h after that, the media containing lentiviral particles were accumulated and filtered through a sterile 0.20 μm syringe filter (Millipore, Germany) to infect target cells. Three days after transduction, infected cells were selected using 2 mg mL^-1^ puromycin (Sigma Aldrich, USA) in the respective culture medium.

#### 2.7.4. Spheroid culture

Q-serts for hanging drop spheroid culture were 3D-printed, processed, sterilized, and used in a 96-well plate as described by Butelmann et al. (2022)^26^. Briefly, to form multicellular spheroids, MDA-MB-231 BFP, HPMEC, and MSC E2 Crimson were mixed in a ratio of 5:3:2, and a 35 µL drop containing 21,875 cells was pipetted through a Q-sert opening. The culture medium was prepared according to the given ratio for the cells. Cells were allowed to settle for 2 days, after which an initial medium change (-5 µL, +7.5 µL) and a subsequent daily medium change (-5 µL, +6 µL) were applied. Spheroids were harvested on day 7 by a short, pulsed centrifugation at 800 rpm at room temperature and collected in a microcentrifuge tube using a pipette after removing the Q-serts.

#### 2.7.5. Imaging and microscopy

Photographic images were taken using an iPhone SE 2020 (Apple, USA). Optical and fluorescence microscopy images of the free and embedded spheroids were obtained with a Zeiss Axio Observer Z1 (Carl Zeiss, Germany).

#### 2.7.6. Software and statistical analysis

For data analysis, Origin Pro 2023 (OriginLab Corporation, USA) and GraphPad Prism 8.4.3 (GraphPad Software, USA) were used. A statistical analysis between only two conditions was carried out using the unpaired t-test in Microsoft Excel (Version 2502, Microsoft, USA). The sample variance was analyzed using the F-test Two Sample for Variances. Analysis of variance (ANOVA) was applied with the Tukey’s test, using OriginPro 2023 (OriginLab Corp., USA). P-values ≤ 0.05 were considered statistically significant. n.s. was used to indicate non-statistically significant results. Number of replicates (n) is indicated for each analysis.

## 3. Results and Discussion

### 3.1. The role of sugars in yeast EPS production

In our previous study, physiological mechanisms that allows *R. toruloides* yeast to produce EPS were identified, indicating that a nitrogen metabolism-dependent acidification triggers EPS biosynthesis^20^. The study showed that EPS production occurs through glycolytic metabolism in the presence of inorganic, but not organic, nitrogen. However, it did not examine the influence of different sugars on EPS production or characterize the properties of the resulting EPS, both of which are relevant for developing applications. Therefore, the present research pursues these scientific questions and leverages EPS for a practical application.

Here, the impact of different glycolytic carbon sources on EPS production was evaluated. Besides glucose (C6), mannose (C6), galactose (C6), or xylose (C5) were selected to cultivate *R. toruloides* in shake flasks (microaerobic condition). The physiology results show that *R. toruloides* could consume both C6 and C5 carbon sources; however, efficiency of carbon utilization and corresponding growth varied (Fig. 1). Similar biomass amounts were achieved for cells grown in media containing glucose (C6) and mannose (C6), reaching DCW of 7.33 ± 0.40 and 8.86 ± 1.00 g L^-1^, respectively, after 72 h of cultivation. In contrast, lower DCW values were observed for galactose (C6) (1.16 ± 0.23 g L^-1^). Xylose (C5) showed an intermediate DCW value (6.15 ± 0.21 g L^-1^) at 72 h, and, differently from the other carbon sources, had a longer lag phase that extended beyond the first 24 h of cultivation, causing the yeast to reach the stationary phase only after 72 h, in this case readouts were collected at 96 h of cultivation, resulting in a DCW of 5.97 ± 0.70 g L^-1^. Those differences could be partially attributed to glucose, xylose, and mannose sharing some of the same facilitated transporters for sugars, while galactose uptake occurs in transporters that are specific for it in yeasts^27^. Moreover, galactose has a complex regulatory network and the expression of its transporter is inducible, in contrast with other hexokinases^28^. Regarding xylose, it is suggested that *R. toruloides* has specific xylose transporters and xylose metabolism efficiency is compromised due to the accumulation of D-arabitol as a byproduct^29^, explaining the limited growth when compared to glucose and mannose. The metabolic pathways connecting each sugar assimilation to glycolysis is shown in Fig. S1-A^22,27,29,30^. Our biomass and sugar utilization results are similar to those reported for *R. toruloides* IFO0880, cultivated on different sugars at 20 g L^-1^ of starting concentrations^27,31^.

**Figure 1.**
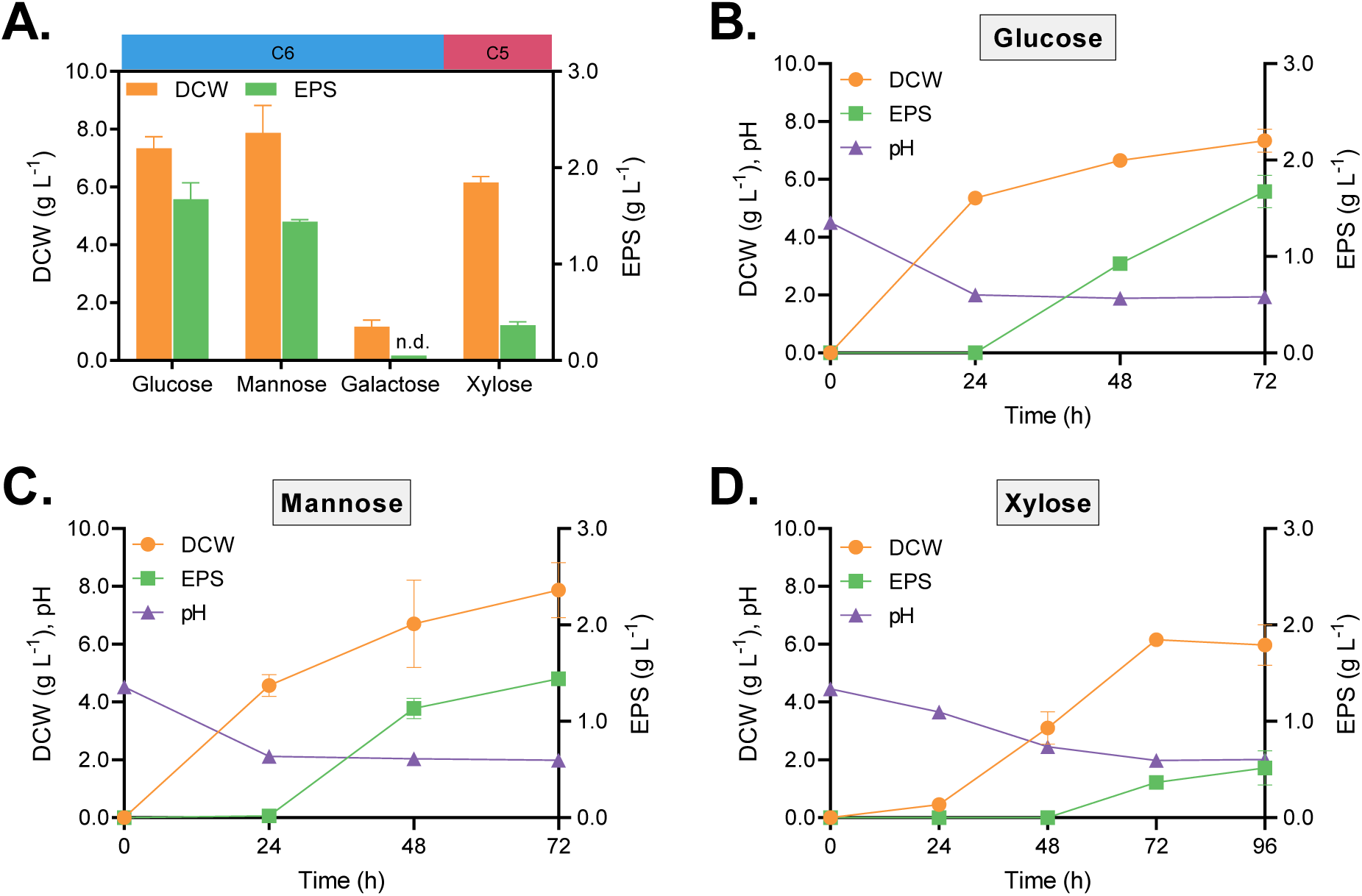
*Rhodotorula toruloides* physiology in shake flasks on different sugars. **A.** Summary of final DCW (g L^-1^, left) and EPS (g L^-1^, right) concentrations. C6 and C5 refer to carbon atoms per sugar molecule. Data corresponds to 72 h of cultivation. Time series cultivation data with DCW (g L^-1^), EPS (g L^-1^), and pH for *R. toruloides* grown on EPS-producing sugars: **B.** glucose; **C.** mannose; and **D.** xylose. Abbreviation: n.d. means EPS was not detected. All experiments were performed in triplicate. Values are presented as mean ± SD.

A carbon source dependency for EPS biosynthesis was observed for all investigated sugars, except for galactose-containing media, where EPS was not detected (Fig. 1-A). Culture acidification was also identified in producing conditions (Fig. 1-B,C,D), consistent with our previous study^20^. EPS production increased over time and amounts were dependent on the carbon source. At 24 h, the pH of the media with glucose and mannose decreased to 2, while the xylose-grown cultures remained at pH 3.65 ± 0.02. Later, at 48 h, EPS was obtained for glucose and mannose media and increased until 72 h, reaching 1.68 ± 0.17 g L^-1^ and 1.44 ± 0.01 g L^-1^, respectively. EPS was first observed in the xylose cultivation at the 72-h time point (0.37 ± 0.04 g L^-1^), when the pH decreased to 1.97 ± 0.01, and further increased at 96 h (0.48 ± 0.12 g L^-1^). The growth profile for the *R. toruloides* is shown in Fig. S1-B. Our results show the role of different C6 and a C5 sugar on EPS yield, which has previously been reported for other *Rhodotorula* metabolites, including lipids, carotenoids, and alcohol-sugars^32,33^. Interestingly, sugars utilization, except galactose, shows similar acidification patterns of *R. toruloides* cultures, unlike the differences observed between inorganic and organic nitrogen utilization in the previous study^20^.

The DCW and EPS results allow determination of overall EPS yields from biomass (Y_PX_, Equation 1) for each sugar, enabling quantification of biomass-normalized EPS synthesis. Under the conditions tested, cells grown in glucose-containing media showed a higher Y_PX_ (0.224 ± 0.008 gEPS gDCW^-1^) after 72 h of cultivation compared to mannose (0.173 ± 0.011 gEPS gDCW^-1^) and xylose (0.059 ± 0.004 gEPS gDCW^-1^). At 96 h, the Y_PX_ for xylose further increased to 0.085 ± 0.019 gEPS gDCW^-1^, likely due to a late EPS production induced by slower acidification of the media. Our results are consistent with previous reports obtained using *Rhodotorula* and *Rhodosporidium* species, where sucrose and glucose are the sugars most frequently used to yield the highest EPS amount^34,35^.

Together, our results suggest that the tested sugars are assimilated by *R. toruloides* through differently regulated sugar transporters, converging on the central carbon metabolism and producing EPS precursors (Fig. 1, S1). Since glucose-grown *R. toruloides* cultures showed highest EPS conversion and Y_PX_, the bioprocess parameters for this condition in batch bioreactors (aerobic) were evaluated.

### 3.2. Bioprocess parameters evaluation for EPS-producing yeast

The bioprocess parameters for EPS-producing *R. toruloides* yeast in 1 L aerated stirred tank bioreactors were evaluated. The aerobic conditions were maintained by keeping the dissolved oxygen level above 50% during cultivation (Fig. S2). All batch experiments were performed in triplicate while monitoring online and offline parameters.

The results show that the profiles of DCW and EPS over 120 h followed a similar pattern as the results presented in Fig. 1-B, in which, by 24 h of cultivation, the yeast approached the stationary phase and started EPS production in shake flasks. However, the DCW continued to slowly increase throughout the cultivation in the bioreactors, increasing from 4.71 ± 0.47 g L^-1^ to 5.38 ± 0.63 g L^-1^, from 72 to 120 h of cultivation (Fig. 2-A). Less biomass accumulation was achieved in bioreactors, since cultures in shake flasks had higher DCW at the end of cultivation, possibly due to differences in parameters between bioreactor and shaker cultivations, including mixing and air flow, allowing rapid acidification and slowing growth. In bioreactor cultivations, two EPS precipitates were obtained after the ethanol precipitation step. The first fraction floated at the top (Top fraction) of the mixture of ethanol and supernatant (2:1 in volume) after 24 h, while the second fraction sedimented to the bottom (Bottom fraction). This dual fractionation could be due to the polysaccharide chain collapse in the bottom EPS^36^. For titer quantification, the EPS fractions were considered together, resulting in 2.67 ± 0.45 g L^-1^ of EPS after 120 h of cultivation. The specific EPS production rate (rEPS, gEPS gDCW^-1^ h^-1^) was calculated following section 2.5, and the results show a spike in EPS production at 48 h of cultivation, followed by a decrease in rEPS for the next points (Fig. 2-B). Regarding off-gas and pH data (Fig. 2-C), the concentrations of CO_2_ and O_2_ varied heavily, relative to their baseline, during the first 24 h of cultivation. This phenomenon is related to the exponential phase of yeast growth, during which the cells consume O_2_ and release CO_2_ at higher rates. Simultaneously, the start of the stationary phase occurred when the pH approached 2.0 at 20 to 24 h of cultivation, which implies that environmental acidification may be one of the factors that interrupted the exponential phase. Our previous study has highlighted the regulatory effects of low pH values in EPS production by *R. toruloides*^20^. Similar results were also reported for *Rhodotorula glutinis*^37^.

**Figure 2.**
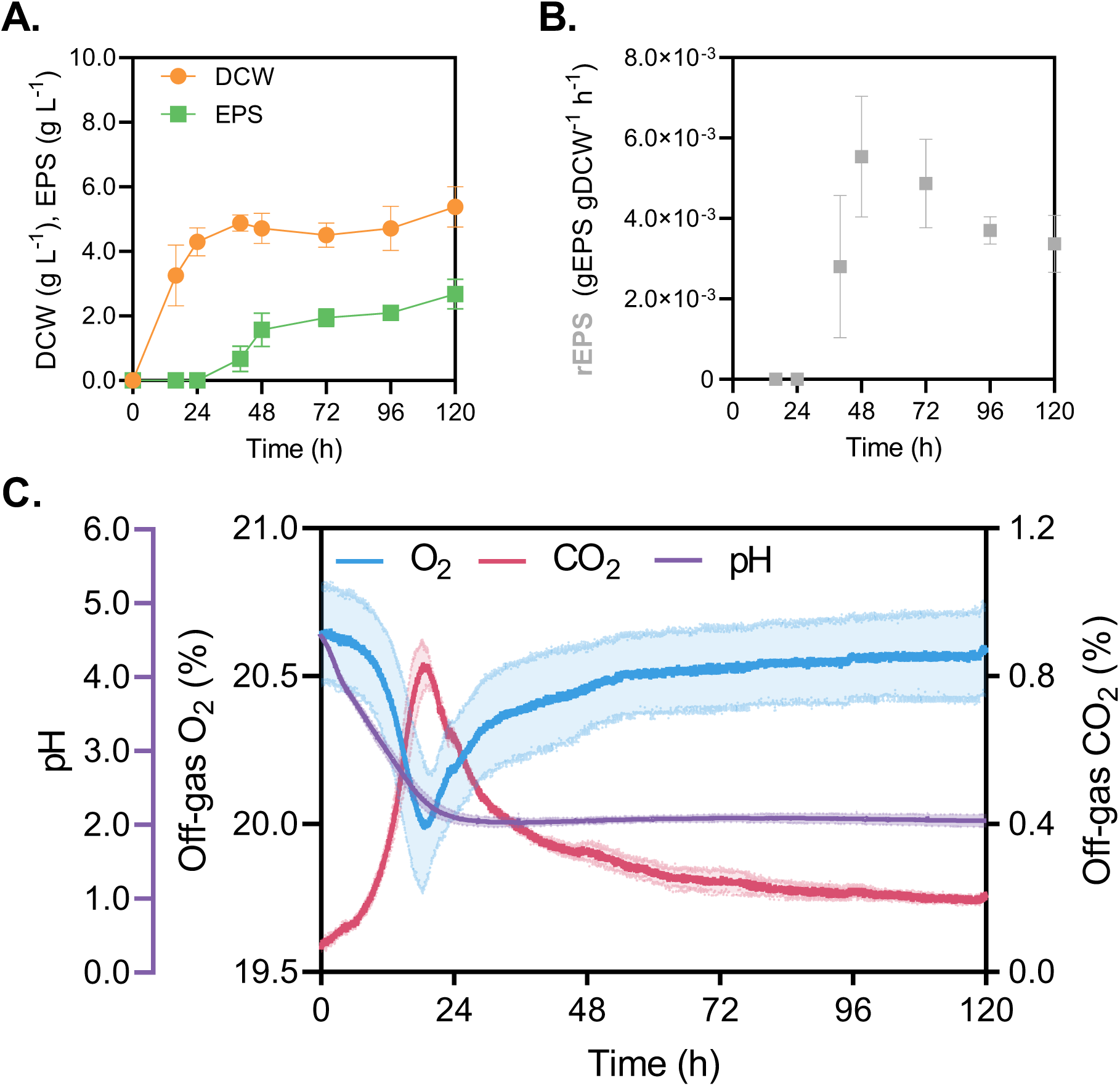
Physiology profiles for *R. toruloides* cultivated in 1 L bioreactors: **A.** DCW (g L^-1^) and EPS (g L^-1^) concentration profiles; **B.** Specific EPS production rates (rEPS) at different time points; **C.** Online bioreactor data for pH, O_2_ off-gas (%), and CO_2_ off-gas (%). All experiments were performed in triplicate. Values are presented as mean ± SD.

The bioprocess parameters of the 1 L batch experiment were determined by first obtaining the maximum specific growth rate (μ_max_) using the CO_2_ production rate (R_CO2_) to identify the exponential growth phase (Fig. 3-A, section 2.5). The exponential phase is observed in the time window between 6 and 15 h after the start of cultivation. The average μ_max_ was 0.2274 ± 0.0010 h^-^_1_ (R^2^ > 0.96). The doubling time for the *R. toruloides* under the studied conditions was 3.05 h. Previous reports show that the μ_max_ largely varies according to the cultivation conditions. González-García et al. explained that for *Rhodosporidium toruloides* ATCC 204091 (previously named as *Rhodotorula glutinis*), media containing ammonium sulfate as the nitrogen source allow for higher μ_max_ values^38^. However, growth in non-buffered media with ammonium sulfate is prematurely stopped due to the decrease in the pH^39^. Lopes et al. demonstrated that μ_max_ also varies depending on the carbon source and C:N ratio utilized, in which a high C:N ratio decreased the μ_max_ for media with acetic acid, xylose, or glycerol as the carbon sources when cultivating *R. toruloides*^33^.

**Figure 3.**
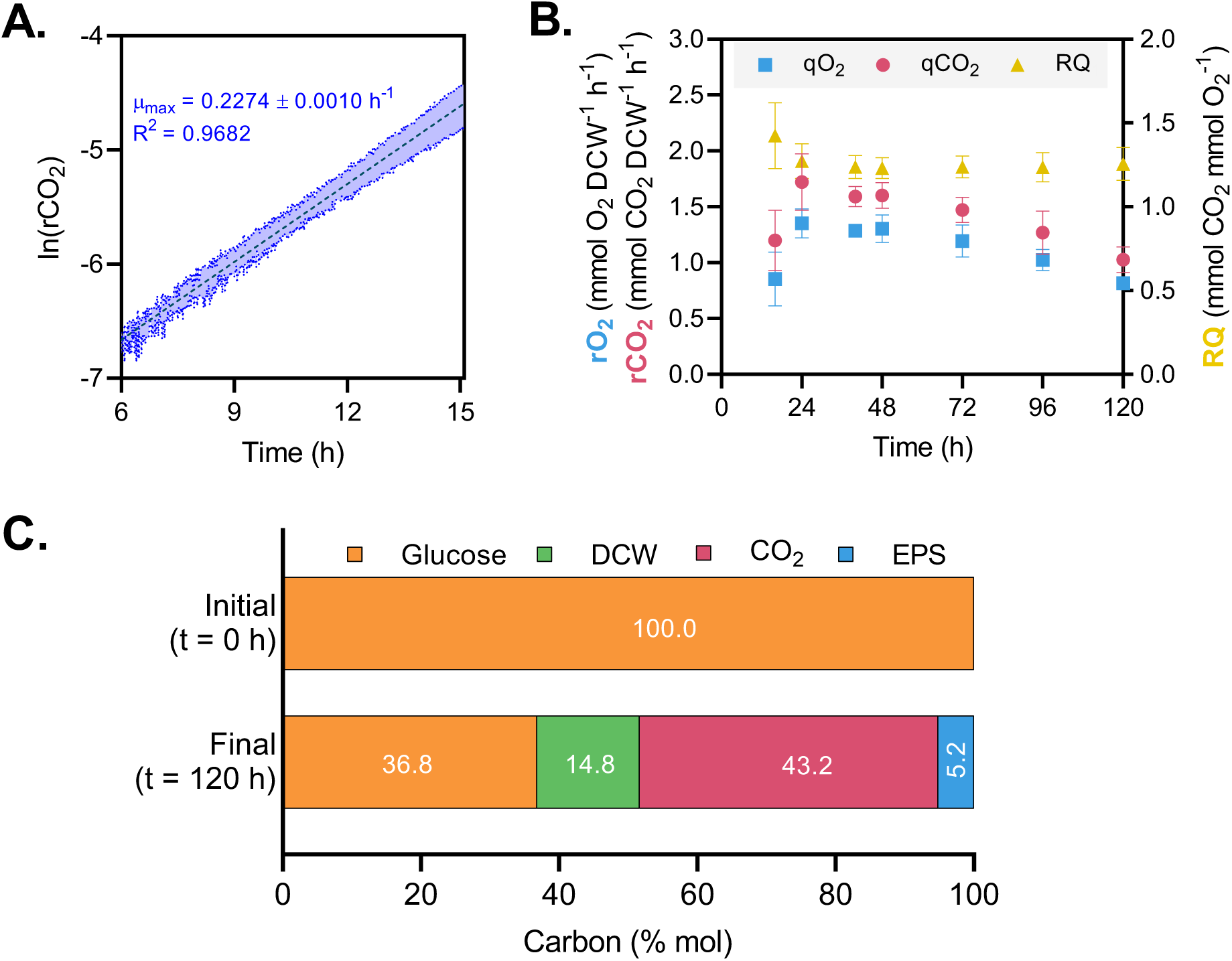
*Rhodotorula toruloides* grown in stirred tank aerated bioreactors: **A.** Average natural logarithm of CO_2_ production rate over time used to determine the maximum specific growth rate (μ_max_). μ_max_ is the slope of the curve. Shaded area is the standard deviation of the replicates; **B.** Specific O_2_ consumption rate (rO_2_), specific CO_2_ production rate (rCO_2_), and respiratory quotient (RQ) over cultivation time; **C.** Carbon balance at 120 h of cultivation. All experiments were performed in triplicate. Values are presented as mean ± SD.

Then, the specific production and consumption rates of CO_2_ (rCO_2_) and O_2_ (rO_2_), and the respiratory quotient (RQ) were calculated (Fig. 3-B) following the Equations in section 2.5. The data used for the calculations are shown in Table S1. The results indicate an increase in both parameters until 24 h, decreasing over time to the initial values, demonstrating a slow reduction in metabolic activity as the process progresses. Moreover, RQ, which represents the relationship between CO_2_ and O_2_, ranged from 1 to 1.5 throughout the experiments, which is reported to occur during lipogenesis or overflow metabolism in oleaginous yeasts, including *R. toruloides*^33,40^.

Furthermore, the carbon balance for the bioreactor cultivation is shown in Fig. 3-C and the complete calculations are on Table S1 and S2. It was assumed that glucose was the only carbon source available at the start of the cultivation, due to the low number of cells inoculated (initial OD_600_ = 0.2). At 120 h, the main carbon-containing products are biomass, CO_2_, and EPS. Glycerol, a minor side product of this yeast^41^, was not detected and was not considered in calculations. The concentration of the remaining glucose obtained by HPLC analysis was 16.83 ± 0.34 g L^-1^, indicating that only approximately 63% of the initial carbon was consumed after 120 h of cultivation, likely due to an inhibition effect caused by the acidification of the medium. This result allows the calculation of the cumulative DCW and EPS yields from substrate (Y_XS_ and Y_PS_, respectively), which were equal to 0.21 ± 0.03 gDCW (g glucose)^-1^ and 0.09 ± 0.01 gEPS (g glucose)^-1^. The minimal biomass composition formula for *R. toruloides* chosen was CH_1.83_O_0.53_N_0.16_ (24.55 g/mol C), based on previously reported data^42^. DCW accounted for 15% of carbon, while CO_2_ production had the biggest share with 43%. Finally, the mass of the monomeric unit was assumed to be 162.15 g mol^-1^ to obtain the EPS’s carbon. Together with the gel permeation chromatography results, shown in section 3.5, EPS was responsible for the remaining 5.2% of the carbon balance. Relative to the consumed glucose, 23.3, 68.4, and 8.3% of the carbon was directed into biomass (DCW), CO_2_, and EPS, respectively.

In summary, when *R. toruloides* was cultivated in a bench scale bioreactor, EPS was efficiently produced in gram-scale amounts. Though growth and carbon uptake were constrained by medium acidification, it triggered EPS biosynthesis in *R. toruloides*. The determination of bioprocess parameters enabled a better understanding of *R. toruloides* metabolism, which would allow EPS production optimization in the future, including in other bioreactor configurations.

### 3.3. EPS composition dependency on different sugars

After evaluating the role of different sugars and the bioprocess parameters for EPS production, chemical and structural characterizations of the extracted EPS were performed, using FTIR analysis for functional groups determination, GC-MS analysis for determining monosaccharide composition, and the Bradford assay for protein quantification.

The analysis of the FTIR spectra of EPS obtained from glucose, mannose, and xylose reveals a typical profile for microbial exopolysaccharides^20,35,43^ (Fig. 4-A). Namely, there is a broad peak for O-H stretching (3226 cm^-1^), double peaks for aliphatic C-H stretching (2926 and 2879 cm^-1^), and for ether CH_2_-O-CH_2_ (1055 and 1016 cm^-1^). EPS chains may contain C=O or bound water due to the presence of peaks at 1674, 1429, and 1375 cm^-1^. The 871 and 743 cm^-1^ peaks show that the EPS has α- and β-glycosidic bonds^44^.

**Figure 4.**
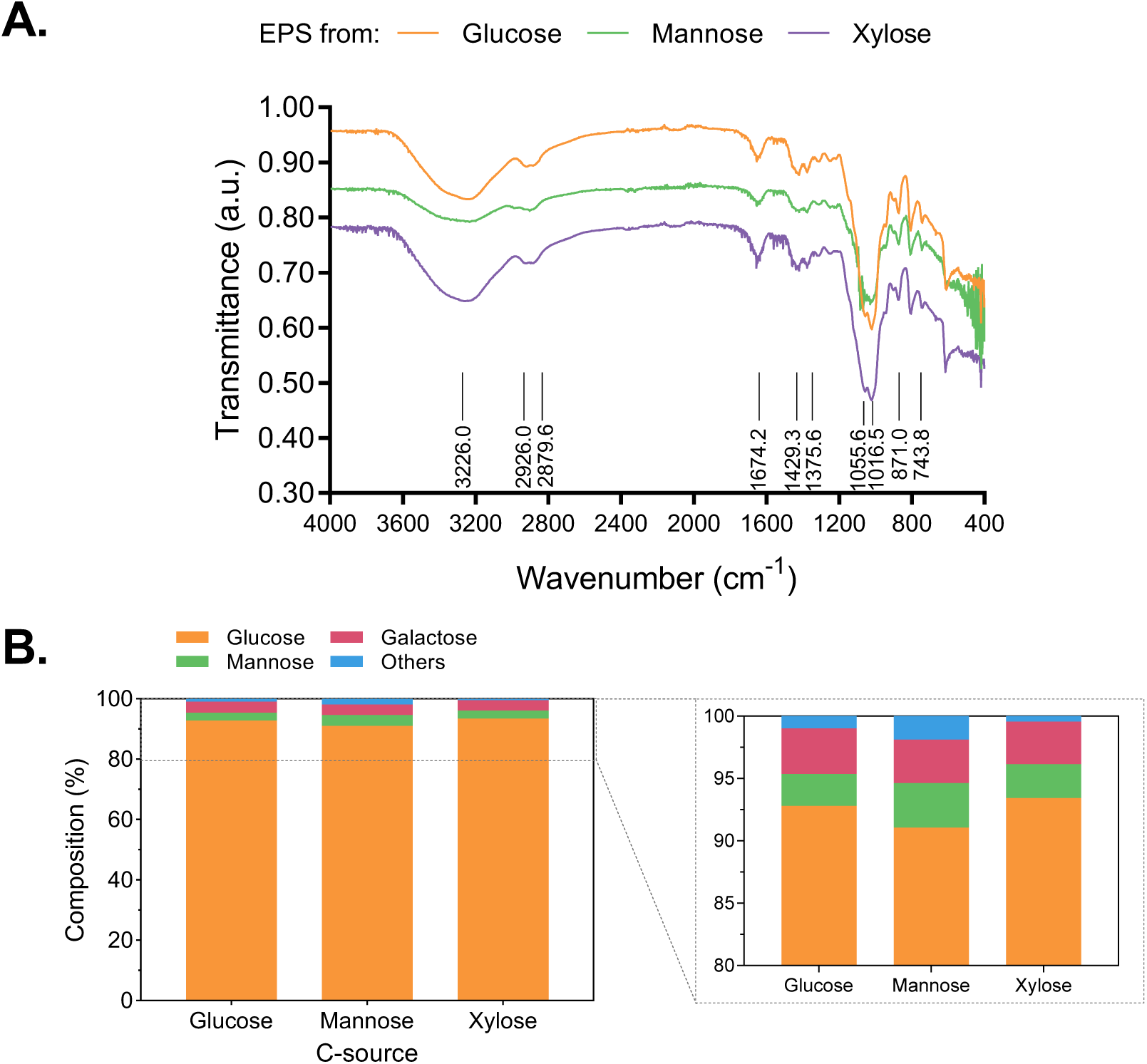
Chemical characterization of EPS obtained from cultivating *R. toruloides* on different sugars. **A.** FTIR spectra for oven-dried EPS obtained from glucose, mannose, and xylose as the only carbon source; **B.** Monosaccharide composition (%) of the EPS grown in glucose, mannose, and xylose as carbon sources. Samples contained glucose 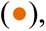 mannose 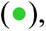 galactose 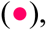 and others 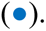 Others include arabinose, xylose, and ribose. All experiments were performed in triplicate.

The monosaccharide composition of EPS slightly varies with the sugars used for its production (Fig. 4-B). The repeating units of EPS were identified by acid hydrolysis of the carbohydrate, followed by the reduction and acetylation of monomeric sugars. EPS derived from glucose, which was also described in a previous publication^20^, was reported to have 92.8 ± 2.9, 2.6 ± 1.3, and 3.7 ± 1.5 % of glucose, mannose, and galactose, respectively. Other monosaccharides, including xylose and arabinose, accounted for 1.0 ± 0.6 %. Similarly, the EPS obtained from yeast cultivated with mannose and xylose sugars had a structure predominantly composed of glucose, with 91.1 ± 2.4% for mannose and 93.4 ± 2.7% for xylose, followed by smaller contributions from mannose (5.3 ± 1.5%, and 2.7 ± 1.1%) and galactose (3.5 ± 1.0%, and 3.4 ± 2.2%) monomers. Arabinose, xylose, and ribose were also detected in samples obtained from cultivations with mannose and xylose, accounting for a total of 1.93 ± 0.7% and 0.6 ± 0.1%, respectively. The relative amounts of the different monomeric units showed insignificant differences, based on each sugar’s triplicate EPS samples. The similarity between the monosaccharide profiles is likely due to a common EPS biosynthesis pathway in this yeast, which our previous study mapped, indicating that the central carbon metabolism supplies the precursors for EPS assembly^20^.

To determine the protein content in the extracted EPS obtained from glucose-containing media, the Bradford method was used, and crude EPS solutions had their absorbance measured at 595 nm. The results show that the protein amount varied from 0.7 to 1.0 wt./vol.%, indicating the EPS had a low amount of protein impurities, likely originating from the biomass.

Overall, EPS with almost identical compositional characteristics are obtained even when *R. toruloides* is cultivated on different carbon sources, confirming an interconnected pathway that generates EPS precursors and mapped in our previous study^20^. Next, due to similar compositional characteristics of EPS from different carbon sources, glucose derived EPS from our bioreactor experiments was used for the determination of glycosidic linkages, and subsequently for molecular weight quantification.

### 3.4. Determination of glycosidic bonds in yeast EPS

To identify the glycosidic bonds present in EPS, it was methylated with iodomethane, then hydrolyzed, reduced, and acetylated, resulting in PMAAs^24^. The summary of glycosidic linkages in EPS samples is shown in Table 1. Moreover, Table S3 compiles the PMAAs information with retention time and mass fragments (m/z) for a more detailed identification. Based on the available information, the backbone is proposed to consist mainly of (1→4)-linked β-glucopyranose (Glc*p*), with (1→3)-linked α-Glc*p*, (2→6)-linked β-Mannopyranose (Man*p*), and β-Galactopyranose (Gal*p*)-(1→6) linkages. The small amount of 3,6-β-Man*p* residues (0.80%) allow branching points in the main chain. The peak area proportions obtained in this analysis were similar to the results from our monosaccharide composition analysis (Fig. 4). Additionally, 42% of the area represented terminal Glc*p*, possibly indicating that side chains terminate in a single residue (Fig. 5). Although some conclusions regarding EPS structure can be drawn through the performed analysis, the complete elucidation of EPS requires further processes, such as ^1^H and ^13^C NMR to increase accuracy on the type of glycosidic linkages through anomeric proton and carbon signal analyses, XRD spectroscopy can be employed to check the degree of crystallinity of the EPS, and the Congo red staining could be used to verify if the EPS chain conformation is a triple helical structure^45^.

**Figure 5.**
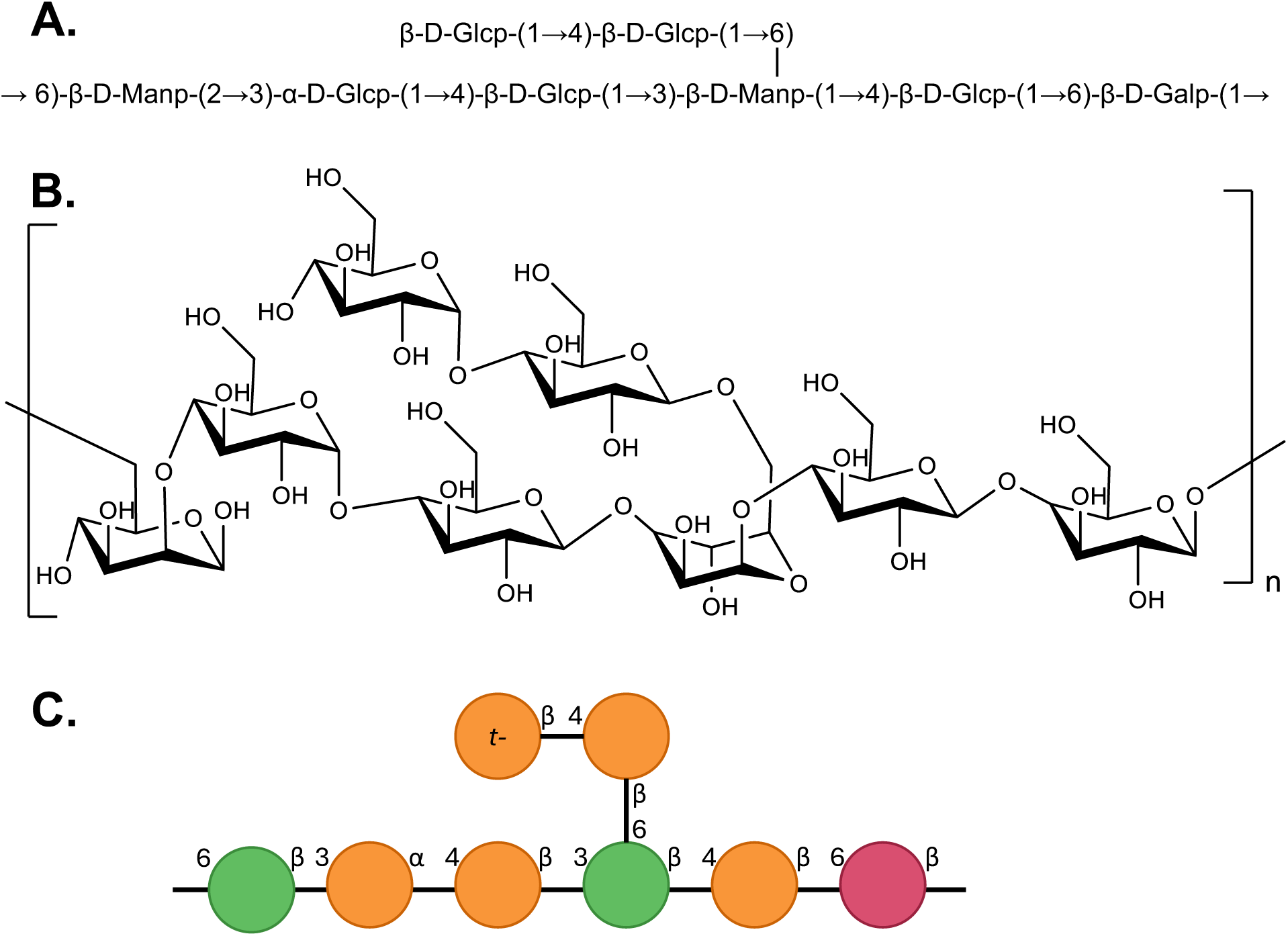
*R. toruloides* EPS putative structure, indicating the identified linkages for the most abundant components: **A.** Standard notation; **B and C.** Schematic representations of EPS structural conformation. Notation: Glc*p* 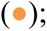 Man*p* 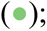 Gal*p* 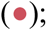 and terminal Glc*p* (t-).

**Table 1.** Glycosidic linkage analysis of *Rhodotorula toruloides* biosynthesized EPS. Percentage area values are presented as mean ± SD of triplicate experiments.

| Partially methylated alditol acetates (PMAAs) | % Area | Linkage type |
| --- | --- | --- |
| Methyl 3-O-acetyl-2,4,6-tri-O-methyl-α-D-glucopyranoside | 7.88 ± 2.10 | → 3-α-D-Glup-1 → |
| 2,3,4,6-tetra-O-methyl-D-glucitol diacetate | 41.61 ± 12.97 | β-D-Glup-1 → |
| 1,3,4-tri-O-methyl-D-mannitol triacetate | 8.29 ± 0.13 | → 6-β-D-Manp-2 → |
| 2,3,6-tri-O-methyl-D-glucitol triacetate | 34.09 ± 5.57 | → 4-β-D-Glup-1 → |
| 2,3,4-tri-O-methyl-D-galactitol triacetate | 7.33 ± 5.17 | → 6-β-D-Galp-1 → |
| Methyl 3,6-di-O-acetyl-2,4-di-O-methyl- α-D-mannopyranoside | 0.80 ± 0.52 | → 3,6-β-D-Manp-1 → |

Methylation analysis has been used to determine the glycosidic linkages in polysaccharide samples, as reported for materials obtained from various microorganisms, including bacteria and yeast. For instance, the EPS from *R. mucilaginosa* GUMS16 was characterized to be majorly branched in both glucose and mannose residues with linkages such as [→3,6)-Glc*p*-(1→], [→2,6)-Glc*p*-(1→], and [→4,6)-Man*p*-(1→]^24^. The EPS from *Zygosaccharomyces rouxii*, which was studied for its cryoprotective ability, was reported to contain mainly [t-Man*p*-(1→], [→2)-Glc*p*-(1] → and [→2,6)-Man*p*-(1→]^46^.

These comparisons indicate that different yeast strains and species have different arrangements of sugar monomers in their EPS structures. As a result, EPS exhibits high structural variability, properties, and opportunities for developing distinct EPS-based applications, highlighting the relevance of our investigation.

### 3.5. Determination of molecular weights range in EPS precipitates

Since molecular weight is a crucial factor that affects polymer applications, GPC was used to determine the molecular weight range for the extracted EPS. For this, EPS from the bioreactor fermentation was employed. The ethanol precipitation method allowed the formation of two distinct EPS precipitates, floating at the top or forming at the bottom of the mixture, as previously mentioned, likely due to a chain collapse on the bottom part (Fig. S3). Both were processed separately and analyzed by GPC to identify any differences in their molecular weight distribution. The top as well as the bottom EPS showed three distinguishable fractions, indicating a high polydispersity index in fractions (Fig. 6-A): fraction 1 with an M_w_ ∼ 1000 kDa, fraction 2 with ∼ 30.0 kDa, and fraction 3 with ∼ 1.75 kDa (Fig. 6-B, Table S4). The areas under the chromatogram peaks show different distributions of the fractions in the two samples, with a shift towards small M_w_ fractions in the bottom EPS (Fig. 6B). Generally, top EPS fractions were more homogenous with a low standard deviation (top EPS max. 3%, and bottom EPS max. 23%). The area of fraction 2 also differs significantly in top and bottom EPS (Fig. 6). It should be noted that fraction 1 possibly consists of two or more sub-fractions due to the identifiable shoulder in the main peak in this fraction (Fig. 6-A).

**Figure 6.**
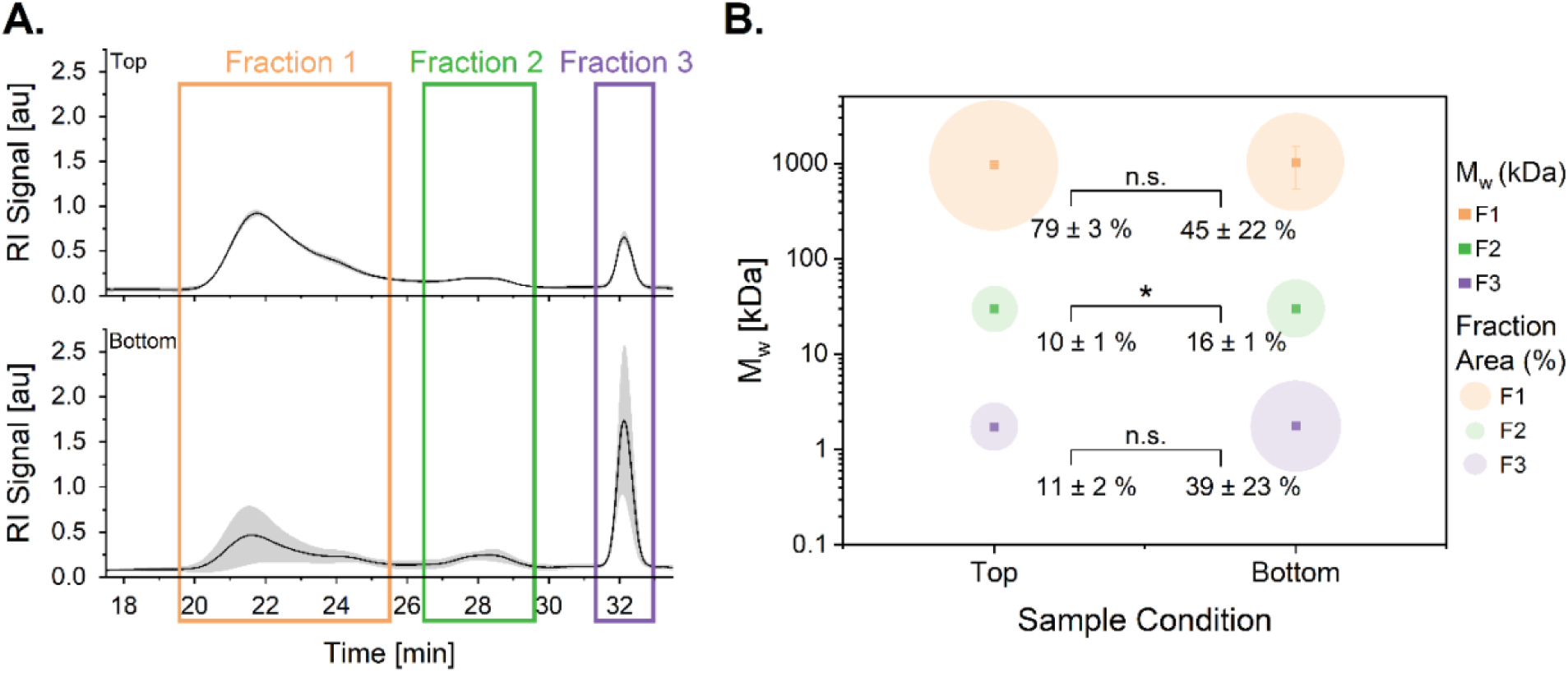
GPC analysis of EPS isolated from top and bottom of the precipitation solution. **A.** GPC chromatograms of top and bottom EPS replicates (grey areas indicating standard deviations); **B.** Molecular weight of the fractions and their respective fraction area represented by relative circles. All experiments were performed in triplicate. Values are presented as mean ± SD, * p < 0.05: significant differences between fraction areas, n.s.: not significant.

Previous reports have shown a wide range of molecular weights for EPS derived from *Rhodotorula* yeasts. Their variability suggests this parameter is specific to each strain and its cultivation conditions. *Rhodotorula glutinis* KCTC 7989 produced a homogeneous mannan with a molecular weight estimated to be around 100 to 380 kDa^37^. Another strain of *R. glutinis*, isolated from blackberry fruits, had a molecular weight of 110 kDa^47^. Two fractions of a mannan EPS from *Rhodotorula acheniorum* MC were characterized to have molecular weights of 249 and 310 kDa^48^. A galactose-rich EPS from *Rhodotorula mucilaginosa* CICC 33013 was reported to have a molecular weight of 7125 kDa^49^.

Generally, in GPC, an interaction of the analyte with the stationary phase should be excluded to have a valid molecular weight. Due to the possible ionic nature of the EPS^20^, it must be considered that the reported M_w_ might differ. However, the measured M_w_ are all in the range of previously reported fungal EPS^50,51^. Static or dynamic light scattering, absolute methods to determine the M_w_ and particle size, can be used to confirm the M_w_ and size of the EPS in the future. Moreover, future studies can potentially focus on isolating and analyzing the different fractions for in-depth characterization with the methodology presented in this study.

### 3.6. Rheological properties of yeast EPS solutions

Natural polysaccharides can have different applications depending on their rheological properties. Therefore, oven-dried EPS was dissolved in ultrapure water to obtain solutions with concentrations ranging from 1 to 30 g L^-1^. Fig. 7 shows that the viscosity of polydisperse EPS solutions increased with higher EPS concentrations. Moreover, the viscosity for all the studied concentrations remained constant between the analyzed shear rate regions, indicating the solution behaved as a Newtonian fluid (Fig. 7-A), likely due to a low density of entanglements from small side chains^52^. This can be an advantage over processes that involve non-Newtonian microbial EPS production processes, where EPS solutions that are highly viscous and shear-thinning limit mixing and aeration, creating heterogenous zones with high and low shear^53^. For instance, bacterial hyaluronic acid production by *Streptococcus zooepidemicus* has limits for maximum yields and titers due to the high viscosity of the cultivation medium when the hyaluronic acid concentration increases^54^. In the future, more concentrated EPS solutions, up to the maximum EPS solubility, could be prepared to verify if the dilution is a factor that governs EPS rheology.

**Figure 7.**
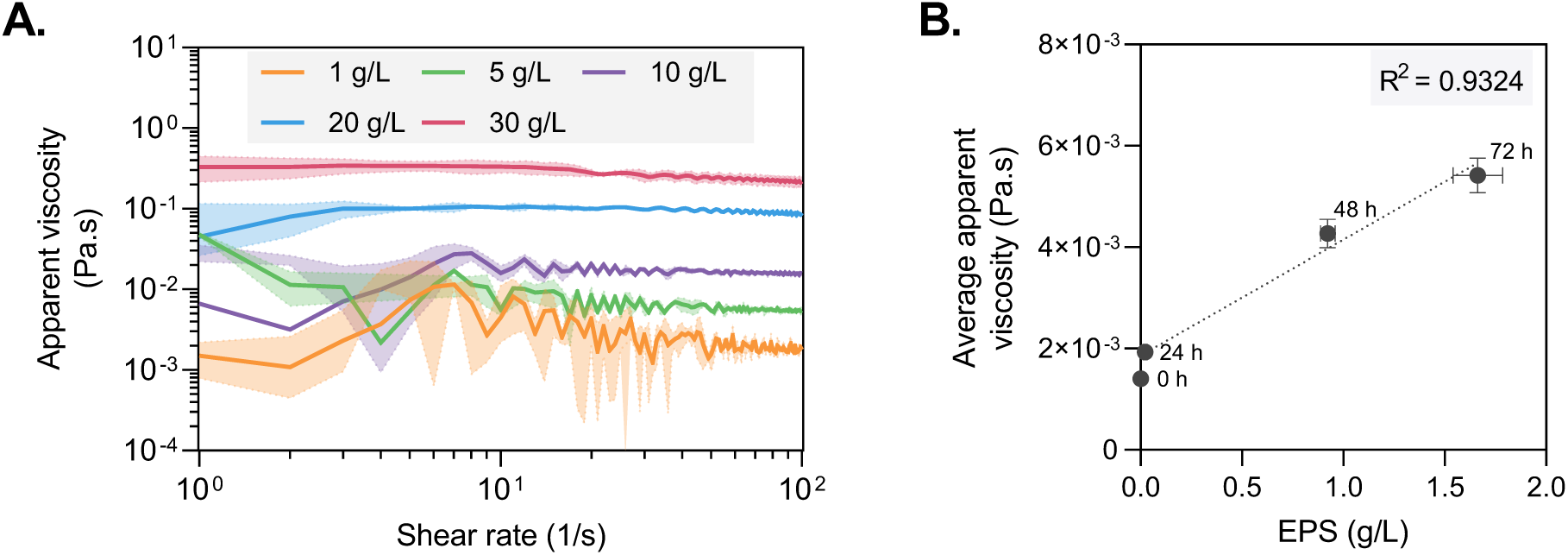
EPS solutions rheology measurements. **A.** Apparent viscosity (Pa.s) over shear rate (1/s) for polydisperse EPS solutions with concentrations: 1, 5, 10, 20, and 30 g L^-1^ (shaded areas indicating standard deviations); and **B.** Average apparent viscosity of culture supernatants over EPS concentration at different time points (0, 24, 48, and 72 h). The initial medium was non-buffered chemically defined medium containing glucose (20 g L^-1^) and ammonium sulfate (5 g L^-1^).

Due to the importance of the viscosity and rheological behavior of the culture broth, the average viscosity of the culture medium in shake flask experiments was correlated with the EPS concentration (Fig. 7-B) for *R. toruloides* grown in chemically defined medium with glucose (20 g L^-1^) and ammonium sulfate (5 g L^-1^). Results show that the average viscosity doubled, going from approximately 2 x 10^-3^ to 4 x 10^-3^ Pa.s, when EPS production was detected at 48 h of cultivation and continued to increase at 72 h when EPS concentration reached 1.68 ± 0.17 g L^-1^, indicating the possibility of monitoring EPS production through the supernatant viscosity. The viscosity measurements over shear rate of the studied supernatant at 0, 24, 48, and 72 h show a near-constant viscosity from 1 to 100 s^-1^ (Fig. S4).

These results indicate that the EPS produced by *R. toruloides* has favorable rheological properties for bioprocessing, minimizing issues that are common in viscous, shear-thinning systems. Finally, the observed properties could favor the applicability of this EPS in developing biomaterials, as shown in the next section.

### 3.7. Hydrogel engineering for embedding and culturing cancer spheroids

To demonstrate biomaterials application of yeast polydisperse EPS, we prepared an EPS-PEGDA semi-IPN hydrogel to embed spheroids for 3D cell cultures (3DCC). We reasoned that microbially produced polysaccharides offer advantages over other naturally sourced alternatives (e.g., alginate) for applications in biomaterials because they can be manufactured with tailorable properties and functionalities by deploying synthetic biology tools^55^. The selection of PEGDA was based on its known uses in tissue engineering, 3DCC, and drug delivery/testing. PEGDA is also regarded as highly cytocompatible, minimally immunogenic, and resistant to protein adsorption^56^. Moreover, since PEG is FDA-approved^57–59^, PEGDA is commonly considered well suited for biomaterial applications. Previously, PEGDA has been combined with microbial polysaccharides in biomaterial design to create or tune key material properties (Table S5).

For hydrogel engineering, we aimed to achieve an EPS solution with a viscosity that would allow proper miscibility of cells, which was accomplished by a 4% EPS (top precipitate) in water. Then, we mixed EPS and PEGDA (4% each, 4% EPS-PEGDA) and performed inversion tests before and after UV-curing in the presence of a photoinitiator (see 2.7.1), comparing the results with proper controls (EPS-only, PEGDA-only, and 4% PEGDA) (Fig. 8-A). In EPS-only solutions, no spontaneous sol-gel transition occurred before or after UV irradiation. For PEGDA-only and 4% PEGDA formulations, we observed a glassy and rigid structure and a matrix that did not fully retain water, respectively. In contrast, the 4% EPS-PEGDA hydrogel remained intact and stable throughout 72 h in cell culture medium, retaining its swelling and water content (Fig. 8-A, Fig. S5).

**Figure 8.**
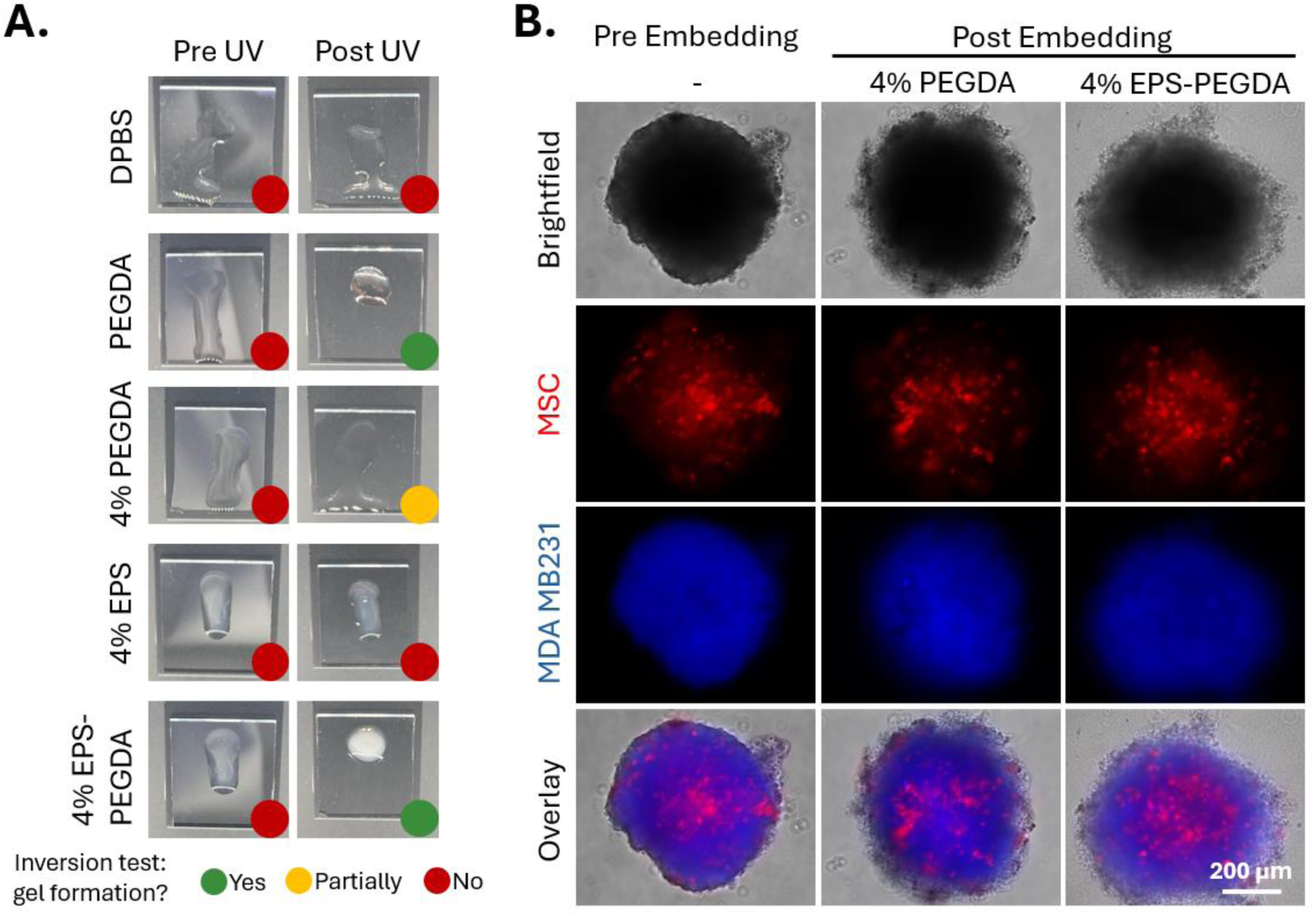
Engineering of an EPS-PEGDA hydrogel for spheroid embedding: **A.** Comparison of different solutions for crosslinking and solidification before and after UV crosslinking; and **B.** Fluorescence microscopy of multicellular spheroids before embedding and after embedding in crosslinked 4% PEGDA, 4% EPS-PEGDA hydrogels.

In previous studies, it has been shown that spheroids consisting of breast cancer cells (MDA MB231), endothelial cells (HPMEC), and mesenchymal stromal cells (MSC) display a vital system for studying the progression of breast cancer *in vitro*^60^. Hence, we used the 4% EPS-PEGDA hydrogel formulation to embed multicellular spheroids composed of the cells described above. The engineered hydrogel enabled fluorescence microscopy imaging of the spheroids, and we qualitatively observed no differences relative to cells embedded in 4% PEGDA (Fig. 8-B).

A systematic evaluation of the 4% EPS-PEGDA and 4% PEGDA hydrogels was conducted to identify differences in qualities that are important for 3DCC applications (Fig. 8 and 9). For both hydrogels, no contamination was detected, and optical as well as fluorescence microscopies were carried out without difficulties (Fig. 9-B). As previously mentioned, in the inversion test, 4% PEGDA did not form a hydrogel that fully retained water, whereas the 4% EPS-PEGDA formulation did (Figs. 8-A and 9-B). Spheroids were retained within the polymeric network after 72 h of cultivation in the 4% EPS-PEGDA replicates, but this was not achieved in the 4% PEGDA (Fig. 9-A, B). As a last point, we evaluated handling by assessing the hydrogel formulation impact on medium replenishment and on the removal of the hydrogels from well plates for endpoint assays and analyses, such as for histological staining. Despite cautious handling, 4% PEGDA was often aspirated and could not be found or easily removed from the well due to its transparency (Fig. 9-A). Conversely, this was easier for 4% EPS-PEGDA, making it the superior choice over 4% PEGDA for reproducible cultures.

**Figure 9.**
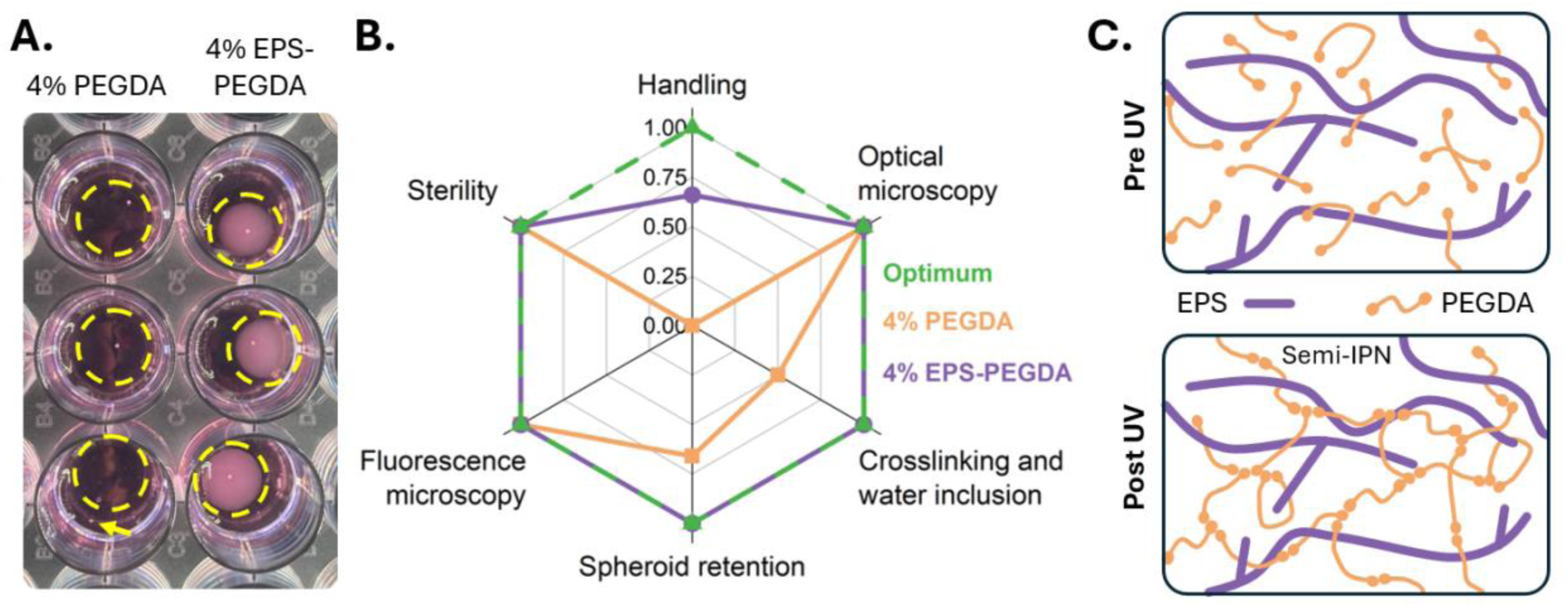
Comparison of hydrogel qualities in cell culture for spheroid embedding: **A.** Three replicates of 4% PEGDA (left) and 4% EPS-PEGDA (right) embedded spheroids in a 24-well plate, dashed circle indicating hydrogel location, arrow indicating spheroid location outside of the hydrogel; **B.** Radar plot comparing performance of both hydrogels, scale in arbitrary units from 0 to 1; **C.** Schematic representation of EPS-PEGDA hydrogel before UV crosslinking and after UV crosslinking as semi-IPN.

The observed changes between the hydrogel formulations could be attributed to both a higher polymer concentration (8% versus 4% total) and the presence of EPS. A key difference is likely the development of a semi-IPN in the EPS-PEGDA hydrogel, where entanglements and inhomogeneities between the PEGDA network and the EPS chains provided new properties and functionalities to the material^61^ (Fig. 9-C). However, more research is needed to fully understand the structure of EPS-PEGDA hydrogels and the mechanisms that govern the interactions between them.

In summary, we have shown the fabrication feasibility of a hydrogel for spheroid embedding using PEGDA and EPS with superior qualities for 3DCC compared to PEGDA-only controls. Our results show proof-of-concept uses of *R. toruloides* EPS in advanced biomaterials. Further modifications and different ratios of PEGDA and EPS could be tested to expand this material to other areas, such as 3D bioprinting and scaffold engineering.

## 4. Conclusions

Our study presents a proof-of-concept biomanufacturing demonstration of yeast EPS to a biomaterial application. Here, we showed the role of various sugars in EPS production, performed EPS chemical and physical characterization, and engineered a hydrogel containing EPS-PEGDA for application in 3DCC, which can be further deployed in potential physiological research models in tissue engineering and biomedicine. In summary, we described the bioproduction conditions and in-depth characterization of the EPS produced by the oleaginous yeast *R. toruloides* CCT 0783, indicating that large-scale production is feasible and can be additionally optimized to enhance substrate-to-EPS conversion and, for example, to enable a multi-product bioprocess by co-producing lipids and carotenoids. We also demonstrated a potential application of this polydisperse EPS by incorporating it with PEGDA in a semi-IPN hydrogel with enhanced biostability and reproducibility compared to PEGDA alone. Future research can be done to expand EPS applications by analyzing the different molecular weight fractions in more detail or chemically modifying the EPS functional groups to make it suitable for specific biotechnological applications. Finally, our study showed that *R. toruloides* yeast has potential for producing extracellular polysaccharides with promising applications in designing advanced biomaterials.

## Supporting Information

**Supplemental 1:** Supplementary Figures (attached file: Supplementary_figures.pdf)

The supplementary figures contain: S1) A. a summarized pathway for consumption of glucose, mannose, galactose, and xylose in *Rhodotorula toruloides*; B. Growth profile of *R. toruloides* cultivated in galactose media; S2) Dissolved O_2_ (%) profiles of bioreactor cultivations of *R. toruloides* over 120 h; S3) Schematic representation of top and bottom EPS fractions after the ethanol precipitation step; S4) Apparent viscosity (Pa.s) of culture supernatants of *R. toruloides* at different time points over shear rate (1/s); S5) Optical microscopy of empty UV-crosslinked controls of 4% PEGDA and 4% EPS-PEGDA over the course of 72 h.

**Supplemental 2:** Supplementary Tables (attached file: Supplementary_tables.xlsx)

The supplementary tables contain: S1) Bioreactor data and bioprocess parameter calculations; S2) Carbon balance summary; S3) Complete information of PMAAs from EPS; S4) Molar mass distribution for different fractions of EPS collected from the top and bottom of the precipitation solution; S5) Literature search of PEGDA combined with biopolymers and their applications as advanced biomaterials.

## Author Information

## Author Contributions

Author’s contribution using CRediT standard: Conceptualization (HSDRH, TB, RK); Data Curation (HSDRH, TB, OT, KO, KLJ); Formal Analysis (HSDRH, TB, OT, KO, KLJ); Funding Acquisition (KO, PJ, VPS, PJL, RK); Investigation (HSDRH, TB, OT, KO, KLJ); Methodology (HSDRH, TB, OT, KO, KLJ, PJ); Project Administration (PJ, VPS, PJL, RK); Resources (VPS, PJL); Supervision (HSDRH, VPS, PJL, RK); Validation (HSDRH, TB, OT, KO, KLJ, PJ); Visualization (HSDRH, TB, OT, KO, KLJ); Writing – Original Draft Preparation (HSDRH, TB, OT, KO, KLJ); Writing – Review & Editing (HSDRH, TB, OT, KO, KLJ, VPJ, PS, PJL, RK). The manuscript was written through contributions of all authors. All authors have given approval to the final version of the manuscript.

## Acknowledgments

The authors wish to thank Ivonne Knauer in the Service Facility for GPC at the Institute for Macromolecular Chemistry at the University of Freiburg. The authors kindly thank Dr. Andrea Barbero for providing the marrow-derived mesenchymal stroma cells. Thanks to Dr. Nicole Gensch from the BIOSS Toolbox, (University of Freiburg, Germany) for providing cells and plasmids.

