## Supplemental Figures for "Engineering Stable Hydrogels with Polydisperse Yeast Exopolysaccharides for Embedding Cancer Spheroids"

**Figure S1** – **A.** Pathways for consumption of glucose, mannose, galactose, and xylose in yeasts, based on *R. toruloides* and *Saccharomyces cerevisiae* literature. HXK1-2: hexokinase I and II; PGI1: phosphoglucose isomerase; PGM: phosphoglucomutase; PMI40: phosphomannose isomerase; XR: xylose reductase; XDH: xylitol dehydrogenase; XK: xylulose kinase; GM: galactose mutarotase; GAL2: galactose permease; GK: galactokinase; URT: uridylyl-transferase. *Means other transporters might be associated with xylose assimilation in *R. toruloides*. Dashed lines mean more than one step is needed to reach the target product. **B.** Time series cultivation data with DCW (g $L^{-1}$), EPS (g $L^{-1}$), and pH for *R. toruloides* grown on galactose.

**A.**

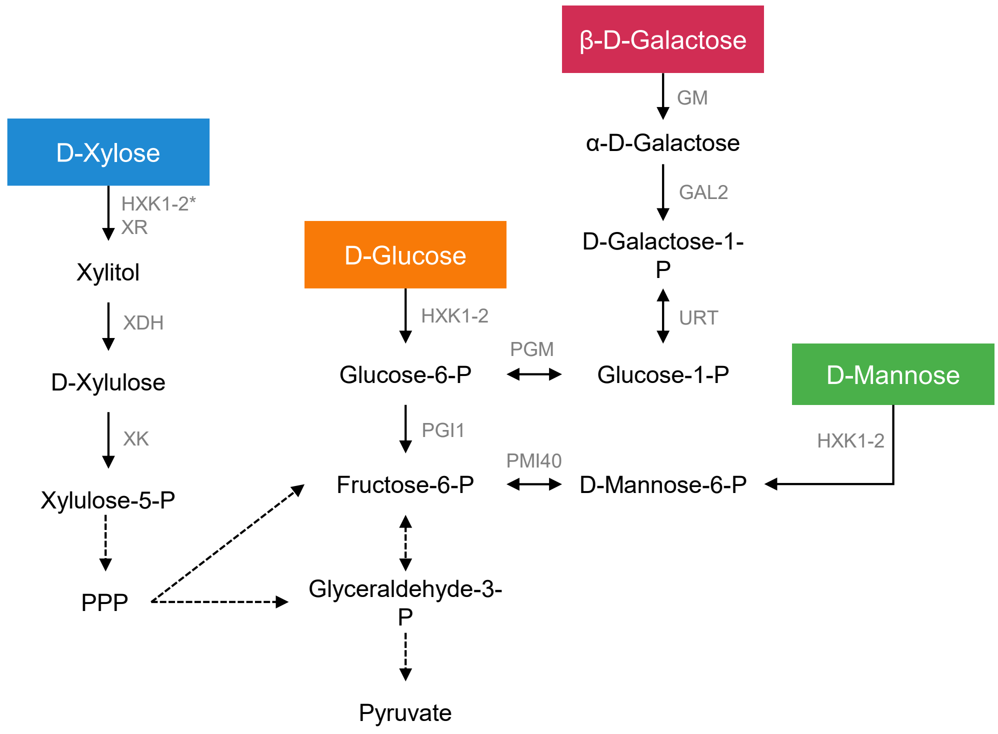


**B.**

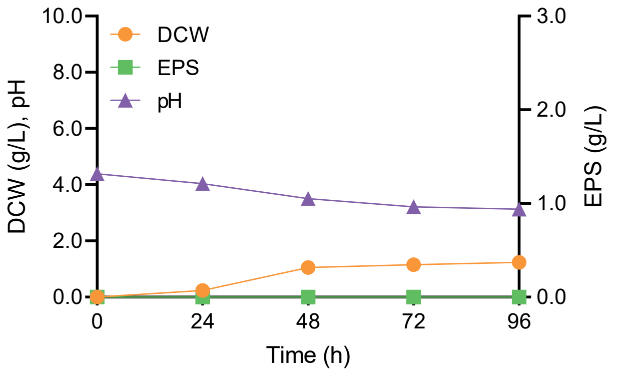

**Figure S2** – Dissolved $O_2$ (%) of bioreactor cultivations of *R. toruloides* over 120 h, indicating that no anaerobic growth occurred. The initial chemically defined medium consisted of glucose (40 g/L), $(NH_4)_2SO_4$ (5 g/L), $KH_2PO_4$ (3 g/L), $MgSO_4 \cdot 7H_2O$ (0.5 g/L), vitamins, and trace elements. The experiment was done in triplicate. Shaded area represents the standard deviation.

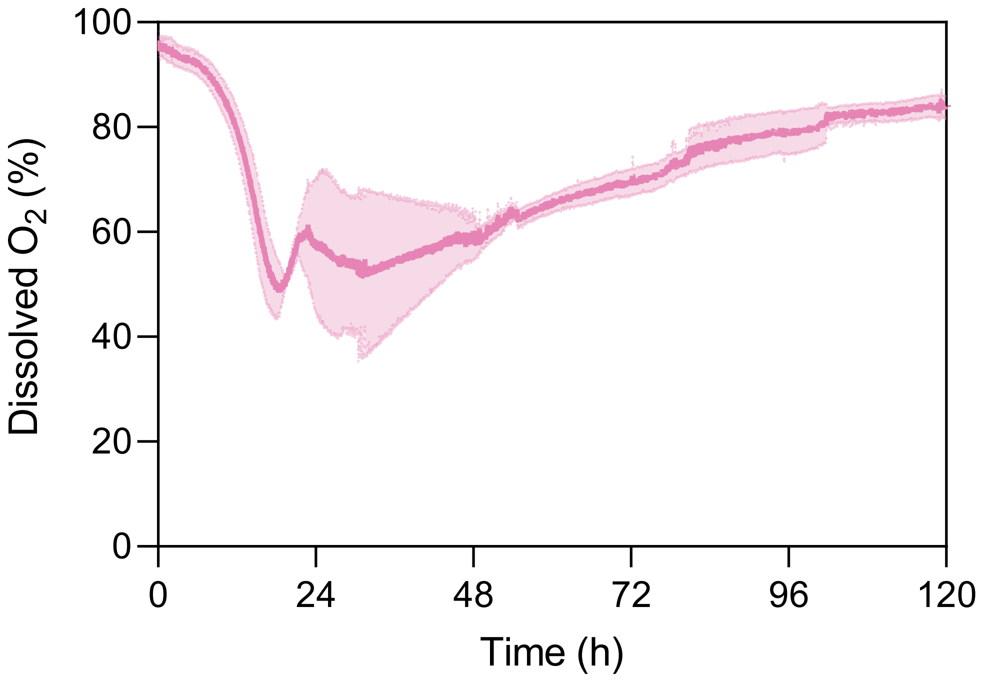

**Figure S3** - Schematic representation of top and bottom EPS parts after the ethanol precipitation step.

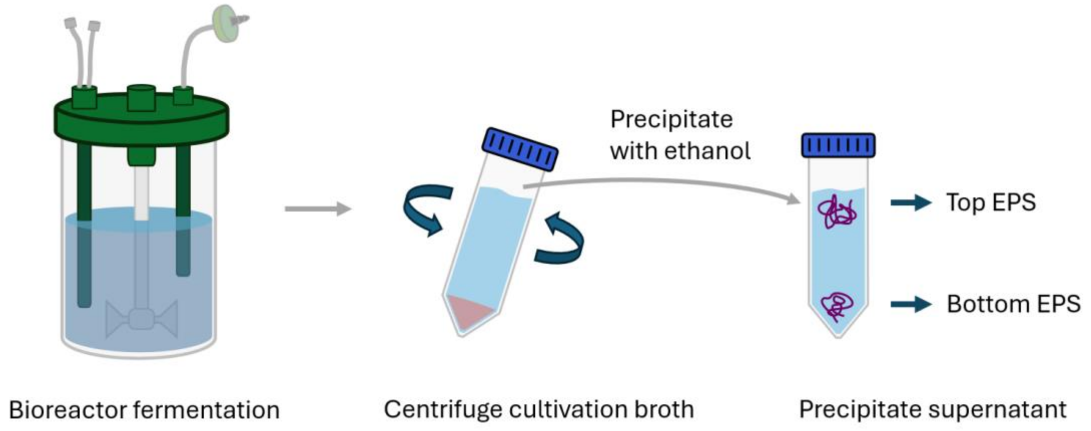

**Figure S4** – Apparent viscosity (Pa.s) of culture supernatants of *R. toruloides* at different time points (0, 24, 48, 72 h) over shear rate (1/s). The initial chemically defined medium consisted of glucose (20 g/L), $(NH_4)_2SO_4$ (5 g/L), $KH_2PO_4$ (3 g/L), $MgSO_4 \cdot 7H_2O$ (0.5 g/L), vitamins, and trace elements.

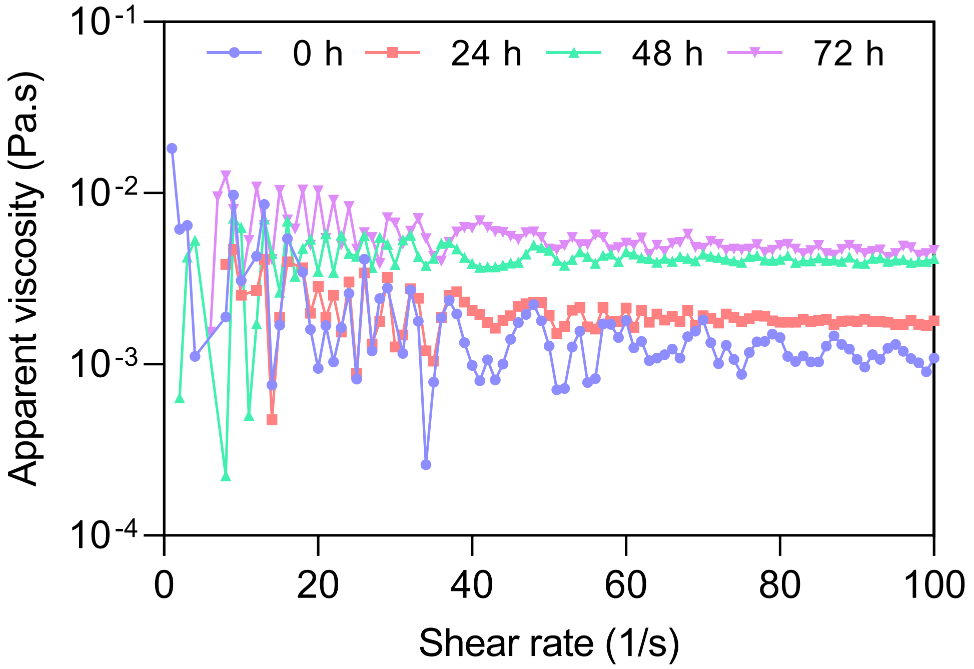

**Figure S5** – Optical microscopy of empty UV-crosslinked controls of 4% PEGDA and 4% EPS-PEGDA over the course of 72 h, hydrogel location left of the observable line.

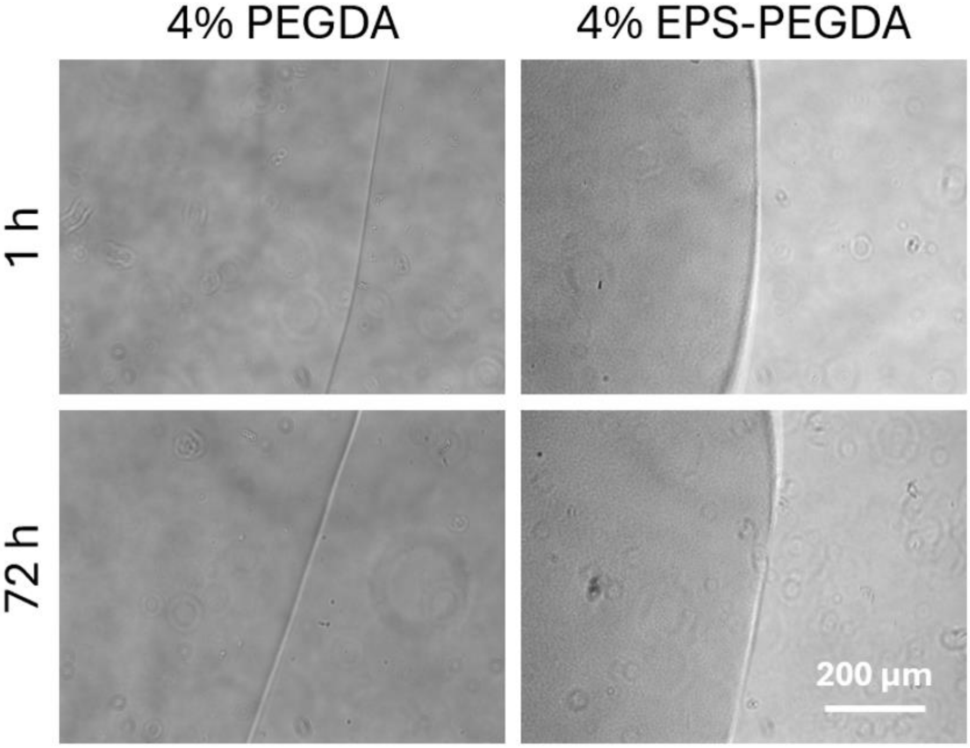
